# FARFAR2: Improved de novo Rosetta prediction of complex global RNA folds

**DOI:** 10.1101/764449

**Authors:** Andrew M. Watkins, Rhiju Das

## Abstract

Methods to predict RNA 3D structures from sequence are needed to understand the exploding number of RNA molecules being discovered across biology. As assessed during community-wide RNA-Puzzles trials, Rosetta’s Fragment Assembly of RNA with Full-Atom Refinement (FARFAR) enables accurate prediction of complex folds, but it remains unclear how much human intervention and experimental guidance is needed to achieve this performance. Here, we present FARFAR2, a protocol integrating recent innovations with updated RNA fragment libraries and helix modeling. In 16 of 21 RNA-Puzzles revisited without experimental data or expert intervention, FARFAR2 recovers structures that are more accurate than the original models submitted by our group and other participants during the RNA-Puzzles trials. In five prospective tests, pre-registered FARFAR2 models for riboswitches and adenovirus VA-I achieved 3–8 Å RMSD accuracies. Finally, we present a server and three large model archives (FARFAR2-Classics, FARFAR2-Motifs, and FARFAR2-Puzzles) to guide future applications and advances.

## Introduction

Noncoding RNA molecules (ncRNA) are essential to biology’s most critical and ancient functions, such as translation (the ribosome), splicing (the spliceosome), and control of gene expression levels (riboswitches) (Cech and Steitz, 2014). Many noncoding RNAs exhibit intricate three-dimensional folded structures, and orders of magnitude more sequences of biologically interesting RNAs have been determined than high-quality RNA structures. For example, there are the thousands of classes of RNA domains that have been curated in the RFAM database but that do not have experimentally solved structures (Kalvari et al., 2018). Therefore, computational methods to predict 3D ncRNA structure could be of substantial value. Current computational methods for 3D ncRNA structure prediction include coarse-grained MD simulation (Vfold3D (Zhao et al., 2017); iFoldRNA (Krokhotin et al., 2015)), coarse-grained Monte Carlo simulation (Rosetta FARFAR (Das and Baker, 2007) and SimRNA (Piatkowski et al., 2016)), and motif assembly (RNAComposer (Popenda et al., 2012); 3dRNA (Jian et al., 2017); MC-Fold (Parisien and Major, 2008)).

Over the last 8 years, these methods have been tested and advanced through the RNA-Puzzles community-wide blind trials (Cruz et al., 2012; Miao et al., 2015, 2017). RNA modeling methods using the Rosetta software (Das and Baker, 2007; Das et al., 2010) motivated the launch of these trials (Sripakdeevong et al., 2012) and have achieved the most accurate models for the majority of cases to date (12 of 21), including ligand-binding riboswitches, ribozymes, and other noncoding RNAs with complex folds. The primary Rosetta modeling tool for RNA, called FARFAR (Fragment Assembly of RNA with Full-Atom Refinement), was inspired by Rosetta’s protein structure prediction methods. FARFAR first models an RNA structure by stitching together 3-residue fragments of previously solved RNA structures whose sequence matches the target sequence. This Monte Carlo process is guided by a low-resolution score function that rewards base pairs and base stacks with geometries similar to those seen in previously solved RNA structures (Das and Baker, 2007). Then, each model is refined in a high-resolution all-atom free energy function that rewards hydrogen bonds, van der Waals packing of atoms, and other physically important interactions, and the lowest energy models are clustered to achieve submitted models (Cruz et al., 2012; Das et al., 2010; Miao et al., 2015, 2017).

While consistently successful in RNA-Puzzles, FARFAR modeling has involved problem-specific expert intuition, such as guesses of ligand binding sites based on inspection of alignments, as well as algorithmic extensions created ‘on-the-fly’ to explore novel ideas inspired by the targets (Cruz et al., 2012; Miao et al., 2015, 2017). As in early days of protein structure prediction (Sali, 1995; Simons et al., 1997), many of these steps have not been well documented, automated, or systematically benchmarked, even as FARFAR has been integrated into practical applications involving experimental data from NMR, chemical mapping, and cryo-EM (Cheng et al., 2015; Kappel et al., 2018; Sripakdeevong et al., 2014). Prior RNA-Puzzles results suggest that some FARFAR-based protocol may potentially have excellent accuracy for modeling complex RNA tertiary folds without experimental data, but the exact protocol and its ability to deliver consistent results remains uncertain due to lack of a systematic benchmark on state-of-the-art code (Ding et al., 2008; Laing and Schlick, 2010; Popenda et al., 2012). Furthermore, without modeling pools associated with such a benchmark, groups developing complementary procedures, including recent scoring methods taking advantage of artificial neural networks and evolutionary coupling information (AlQuraishi, 2019; Li et al., 2019; Wang et al., 2019; Weinreb et al., 2016) and alternative high-resolution refinement procedures (Tan et al., 2018; Watkins et al., 2018), cannot easily test if their procedures might be pipelined with Rosetta FARFAR modeling.

This paper seeks to address these gaps in Rosetta RNA modeling. First, we describe how improvement of the general Rosetta codebase as well as consolidation of the FARFAR protocol have enabled development of a streamlined version of the method, named FARFAR2, which we make available as a webserver on ROSIE at https://rosie.rosettacommons.org/farfar2. The method reproduces and extends prior results on two benchmarks of small RNA folds and RNA submotifs here revisited as the *FARFAR2-Classics* and *FARFAR2-Motifs* model sets. Second, we present benchmarks of FARFAR2 including, most importantly, a new benchmark that revisits every previous RNA-Puzzle for which there exists a final deposited structure. This *FARFAR2-Puzzles* benchmark uses only secondary structure information and template structures employed at the time of the original challenge. Finally, as rigorous validation, we present new blind tests of the method based on four independently solved structures of riboswitch aptamers based on cryo-EM maps (Kappel et al., 2019) and an adenoviral noncoding RNA presented as RNA-Puzzle 24 (Hood et al., 2019). The results confirm the ability of Rosetta FARFAR2 to recover complex global folds of RNAs while also highlighting current limitations of the method that will likely require complementary approaches to resolve. The input files and output models from this study – including over 10 million structures available in a single archive – provide a rich resource that we expect to be valuable in developing approaches that extend or go beyond Rosetta FARFAR2.

## Results

### Consolidated RNA fragment assembly protocol improves modeling of small RNAs

Our core goal in developing the Rosetta FARFAR2 protocol has been to achieve a single application that enables straightforward modeling of complex RNA structures with sizes up to 200 nucleotides, incorporating any available additional knowledge. Previous attempts to construct a comprehensive modeling pipeline required several manual steps running a series of distinct Rosetta applications, such as pre-generating helix ensembles, set up through a separate Python script (Cheng et al., 2015; Watkins et al., 2019). The new FARFAR2 protocol is designed to instead take input information in as simple a manner as possible into a single Rosetta executable, *rna_denovo*. Analogous to other RNA modeling packages (Krokhotin et al., 2015; Piatkowski et al., 2016; Popenda et al., 2012), the *rna_denovo* executable now accepts the RNA sequence, the RNA secondary structure in community-standard dot-parentheses notation, and, if available, the names of PDB-formatted files holding template structures of any known sub-motifs or sub-domains.

FARFAR2 also implements three methodological improvements. It uses an updated library of fragments, based on the non-redundant 2018 crystallographic database of 657 RNA structures (Leontis and Zirbel, 2012), which is 15% larger and more diverse than the previous updated fragment library from 2009 (Richardson et al., 2008). The protocol also implements a special set of Monte Carlo moves for nucleotides in stacked Watson-Crick pairs (“base pair steps”) that maintain Watson-Crick geometry of RNA helices while allowing their backbone conformations to be perturbed, drawing on the same crystallographic database (see Methods). Last, during the minimization stage, the protocol uses an updated all-atom free energy function developed in a recent study seeking high accuracy on small RNA noncanonical motifs (Watkins et al., 2018).

As initial tests of this FARFAR2 protocol, we measured its performance on two benchmarks involving small RNAs developed in prior work. We revisited the original Rosetta RNA benchmark of 18 small RNA problems (Das and Baker, 2007), generating a set of 25.2M total models (1.4M models per problem) that we term the *FARFAR2-Classics* data set. These results confirmed that the new fragment library, minimization score function, and mode of Watson-Crick base pair modeling give improved results over the original Rosetta fragment assembly method, with the most notable improvements arising from full-atom refinement (Figure S1). Full results are given in Table S2. We further assessed performance using the “native-like” standard applied in the original work (Das and Baker, 2007): achieving folds with RMSD to experimental structure of better than 4 Å in the top 5 cluster centers (5000 low-energy models clustered with a 3.0 Å radius). By this metric, FARFAR2 succeeded in 15 of 18 cases, better than the original results of 10 of 18. Figure 2A-E shows native-like models achieved by FARFAR2 for the five cases for which the original study did not sample such folds. For 1A4D (Figure 2A), an NMR structure of the loop D/loop E arm of the E. coli 5S rRNA, the FARFAR2 model correctly recovers eleven consecutive base pairs, only one of which is a canonical Watson-Crick base pair. For 1CSL (Figure 2B), the HIV RRE high affinity site, FARFAR2 recovers an unusual bent geometry and both ‘bulged-out’ nucleotides. For 1I9X (Figure 2C), the branchpoint duplex from U2 snRNA, FARFAR2 recovers a nearly atomic-accuracy model (2.5 Å RMSD). For 1KKA (Figure 2D), an NMR structure of the unmodified anticodon stem-loop from tRNA-Phe, FARFAR2 obtains a model with a correct geometry for the unusually twisted helix, as well as a geometry for the apical loop that lacks several clashes present in the database-deposited coordinates. Finally, for 2A43 (Figure 2E), a pseudoknot from luteovirus, FARFAR2 recovers the A-minor motif that anchors the pseudoknot fold.

**Figure 1.**
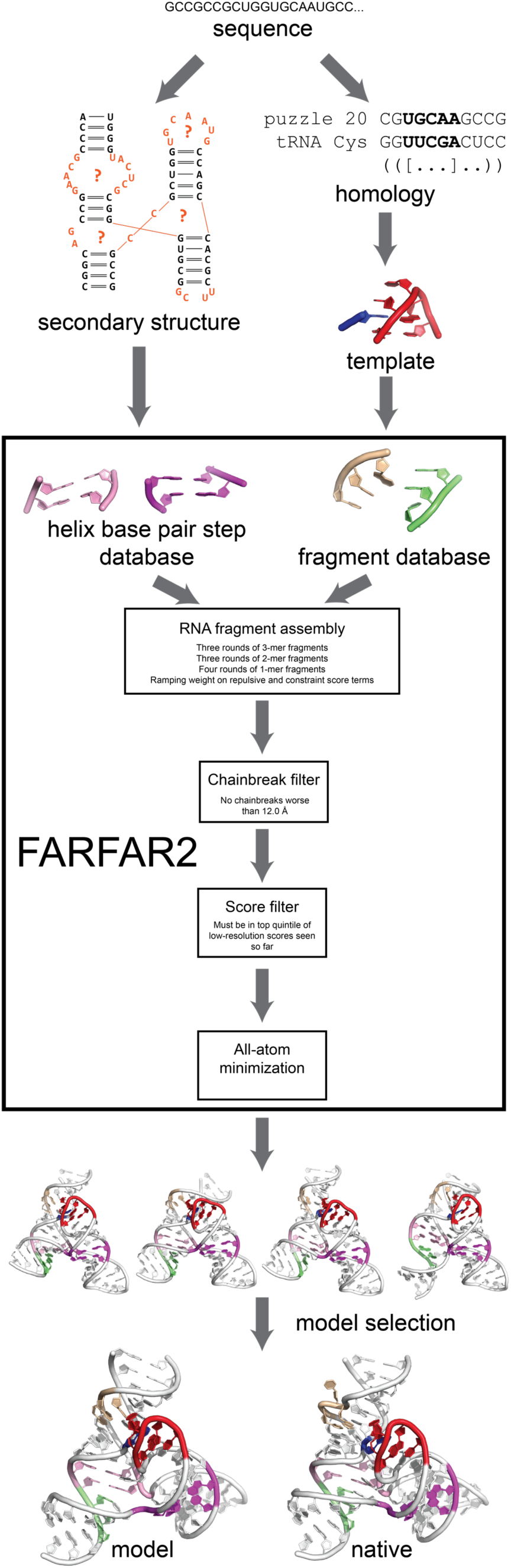
The FARFAR2 structure prediction algorithm. A 3D structure prediction problem is specified by RNA sequence; from that sequence, a consensus secondary structure may be obtained from prior literature studies or covariance analysis of sequence alignments (left), and homologies may be identified to previously solved structures (right). The orange areas in the depicted secondary structure diagram represent the regions whose conformations are unknown *a priori* and whose solution would guide the tertiary structure prediction. Manually identified homologies can also furnish template structures, which are combined by automatic sampling from a base pair step and fragment database in a low-resolution fragment assembly stage. Subsequent models are filtered to omit trajectories with bad chainbreaks or poor scores, and passing models are subjected to minimization in an all-atom scoring function. Finally, models are chosen from the resulting ensemble and compared to the native structure.

**Figure 2.**
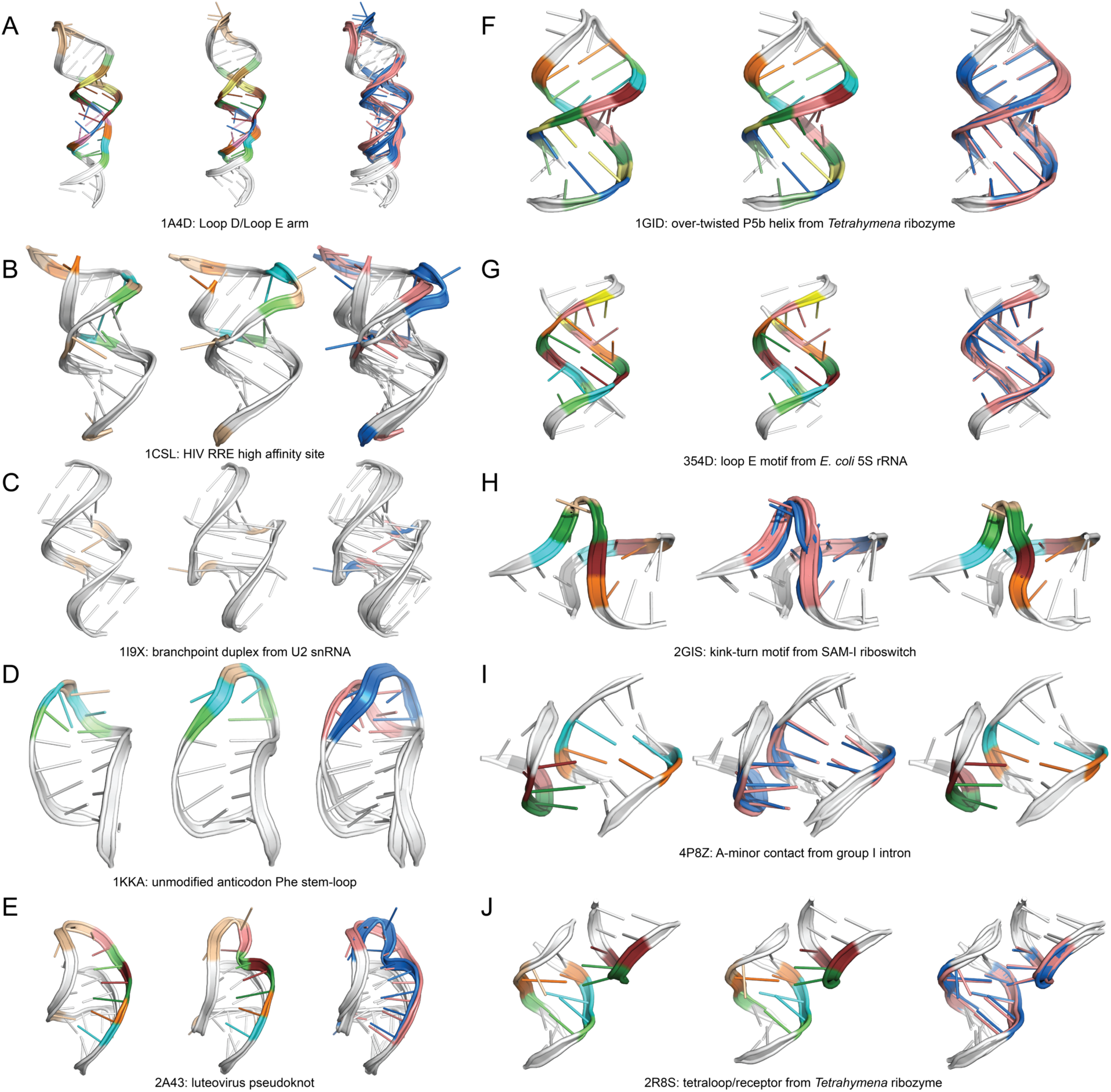
Cases from the FARFAR2-Classics (A-E) and FARFAR2-Motifs (F-I) benchmarks that saw success from the application of FARFAR2 instead of FARNA and SWM, respectively. In each panel, FARFAR2 model, native structure, and overlay are shown from left to right. In (A-E), the FARFAR2 model is the best of 5 low-energy cluster centers after clustering 5000 models with a 3.0 Å radius; in (F-J), the FARFAR2 model is the best of 5 low-energy cluster centers after clustering 400 models with a 2.0 Å radius. In each case, model selection conditions exactly reproduce the conditions used in the original publications using these benchmarks where structures of this quality could not be recovered (Das and Baker, 2007; Watkins et al., 2018). In overlays, the FARFAR2 model is colored in salmon and the native in blue; in individual structures, recovered noncanonical base pairs are colored in cyan, lime, orange, salmon, and ruby, and recovered bulged residues are colored in wheat. In (A-E), residues from pre-specified, flexible helices are colored white; in (F-J), fixed input residues (mostly helical) are colored white.

As a higher resolution test, we evaluated FARFAR2 on a benchmark of noncanonical RNA motifs (apical loops, internal loops, junctions, and tertiary contacts) extracted from larger RNA structures (Das et al., 2010). Recently, we reported that a new nucleotide-by-nucleotide build-up method called stepwise Monte Carlo (SWM) outperformed FARFAR for intricate noncanonical loops (Watkins et al., 2018). However, for problems with longer loops or that require positioning of distinct helical elements in tertiary contacts, the SWM method transited through physically unreasonable intermediate conformations (Watkins et al., 2018), and FARFAR2 achieved better RMSD accuracies than SWM in these cases. Here we revisited this comparison with FARFAR2; the resulting 820,000 models (10,000 models per problem) comprise the *FARFAR2-Motifs* dataset. As observed previously, SWM achieved a 1.5 Å model among the top 5 cluster centers in more problems than FARFAR2 (42 compared to 37 out of 82; see Figure S2 and methods for clustering details). Nevertheless, cases in which FARFAR2 outperformed SWM supported the continuing use of FARFAR2 for modeling complex RNA folds with long loops or tertiary contacts for which the partners’ relative positions are uncertain (Figure 2F-J). An over-twisted helix P5b from the P4-P6 domain of *Tetrahymena* ribozyme, the loop E motif from E. coli 5S rRNA, and the kink-turn motif each involve concomitant modeling of two strands with lengths up to 9 nucleotides, and SWM had difficulty building up complete solutions for these loops (best cluster center RMSDs of 2.7 Å, 1.7 Å, and 2.1 Å, respectively). In contrast, the best of five cluster centers from FARFAR2 did achieve sub-Angstrom recovery of these motifs (0.76 Å, 0.72 Å, and 0.97 Å RMSD, respectively; Figures 2F-H). For tertiary contacts in which the relative positioning of partners had to be modeled *de novo*, current SWM procedures for docking the partner segments gave poor accuracies, e.g., for an A-minor tertiary contact from the lariat-capping GIR1 ribozyme and from the tetraloop-receptor contact of the P4-P6 RNA (1.8 Å and 3.0 Å, respectively). Fragment-based FARFAR2 recovered these structures with excellent accuracies of 1.2 Å and 0.81 Å RMSD, respectively (Figures 2I-J). Because such long-looped junctions and tertiary contacts appear frequently in complex RNA folds, these results have motivated us to continue to develop FARFAR2 as our default procedure for such modeling cases (Watkins et al., 2018, 2019).

### The RNA-puzzles benchmark

The findings above on prediction accuracies for small RNAs and RNA motifs motivated us to test FARFAR2 against larger RNA structures with many motifs and complex folds. Through years of participation in the RNA-puzzles trials (Cruz et al., 2012; Miao et al., 2015, 2017), we have kept records of our strategies for each prediction challenge, including secondary structure predictions, inferences based on homology to prior deposited structures, and functional constraints, such as sites of self-cleavage in ribozyme challenges. We ran FARFAR2 for each of the 21 problems for which an experimental structure is now available. This set comprised all RNA-Puzzles from RNA-Puzzle 1 through RNA-Puzzle 21 except RNA-Puzzle 15 and treated bound and unbound states of RNA-Puzzle 14 (a riboswitch aptamer for glutamine) as separate problems. Consequently, the benchmark included both single- and multi-stranded RNAs, as well as problems for which considerable homology was available and puzzles where we had to start from only secondary structure information (Table S3). We made no explicit provision for ligand binding, save implicitly when the ligand binding site was part of a template structure, since augmenting the low-resolution fragment assembly stage of modeling with a concurrent ligand docking protocol would require substantial additions to the FARFAR2 algorithm.

Our primary question was whether FARFAR2 samples native-like global folds within parts of models of the typical size we would generate for blind modeling challenges. For each of the RNA-Puzzles, we therefore generated 3000-30,000 FARFAR2 models, involving approximately 6-48 hours of computation on 500 CPUs. In typical modeling scenarios, we would inspect approximately 100-200 low energy models as potential candidates for submission, corresponding to the lowest 1% of models by Rosetta all-atom energy. To assess whether any of the large RNA models were native-like, we translated the 4.0 Å RMSD threshold used in our assessment of the smaller RNAs of the FARFAR2-Classics to these new challenges, making use of a previous extension to the length-independent RMSD_100_ metric (Carugo and Pongor, 2008; Kappel and Das, 2019). A 4.0 Å RMSD on FARFAR2-Classics (median length 26 nt, all of which are built *de novo*) is equivalent to a 9.1 Å RMSD on FARFAR2-Puzzles (median length 71 nt that need to be built *de novo*); see STAR Methods. Most problems were close to the median length, but the three longest problems are substantially longer (117, 130, 185 nucleotides must be built for RNA-Puzzles 12, 5, and 7 respectively), and, applying the same transformation, we used 13.8 Å RMSD to assess whether models of these longest problems were native-like.

Using these evaluation criteria, the FARFAR2 protocol is able to sample native-like models within its top 1% of models by predicted energy for 19 of 21 cases. The agreement across all cases is unusually striking, as shown in Figure 3. In 16 of 21 RNA-Puzzles, FARFAR2 samples a model within its best 1% by energy (30–300 models) closer to native than the best originally submitted model during the actual RNA-Puzzles trial (20-100 models; Table 1, Figure 3). In two additional cases, FARFAR2 outperforms any previous Das lab submission but not the very best model overall; detailed depictions of prior models in Figure S3. These results indicate that FARFAR2 emulates or exceeds prior performance in RNA-Puzzles, and specific cases illustrate how this was achieved. In the FARFAR2 model of RNA-Puzzle 6 (Figure 4F), an adenosylcobalamin riboswitch, and of RNA-Puzzle 7 (Figure 4G), the VS ribozyme, the global fold of both RNAs is recapitulated accurately up to one missed inter-helical angle; that level of accuracy is in part enabled by slightly altered helical geometries not found in the original submissions (Figure S4). Marked improvement is also seen in several ribozyme structures of moderate size (Puzzles 15, 17, 19, 20). These molecules – a hammerhead and pistol ribozyme and two bimolecular twister sister constructs – each feature a highly compact multiway junction and a key tertiary contact. The pistol ribozyme features a pseudoknot, while the other three possess intercalated T-loops, and combined with other interconnections, these features lead to slight under- or over-twisting of helices. These observations suggested that the improvements seen in FARFAR2 relative to the original RNA-Puzzles submissions might be due to improvements in helix modeling through “base pair step” fragments. Additional tests using previous helix modeling procedures confirmed the importance of this new helix modeling scheme (Figure S5).

**Figure 3.**
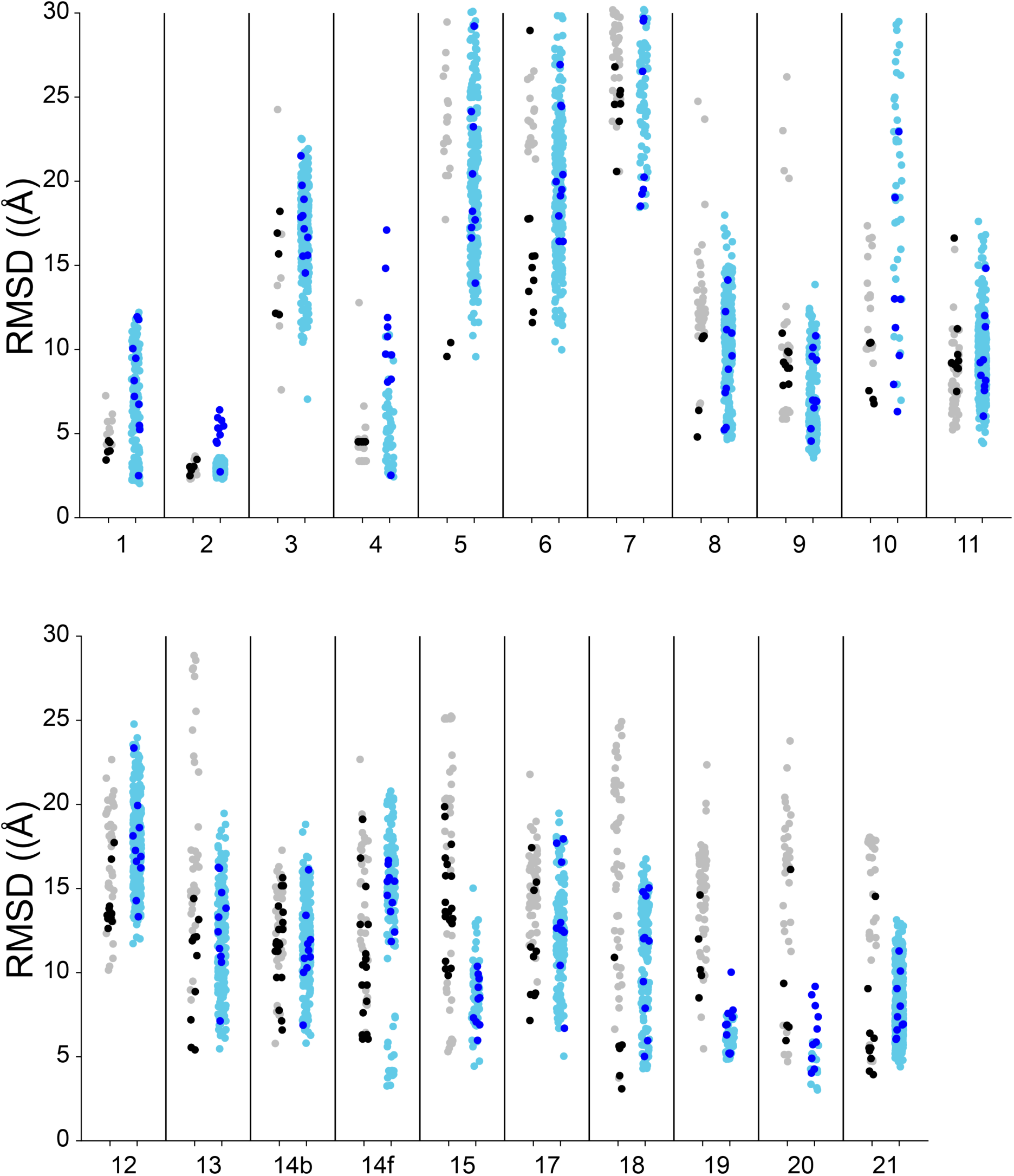
Direct comparison between original RNA-Puzzles submissions from all groups (left points) and FARFAR2 models (right points) for each benchmark case. Among RNA-puzzles submissions, those from the Das lab (created using manually curated Rosetta models, mostly using FARFAR) are black points; others gray. Among FARFAR2 models, light blue points are the top 1% of models by energy; dark blue are cluster centers.

**Figure 4.**
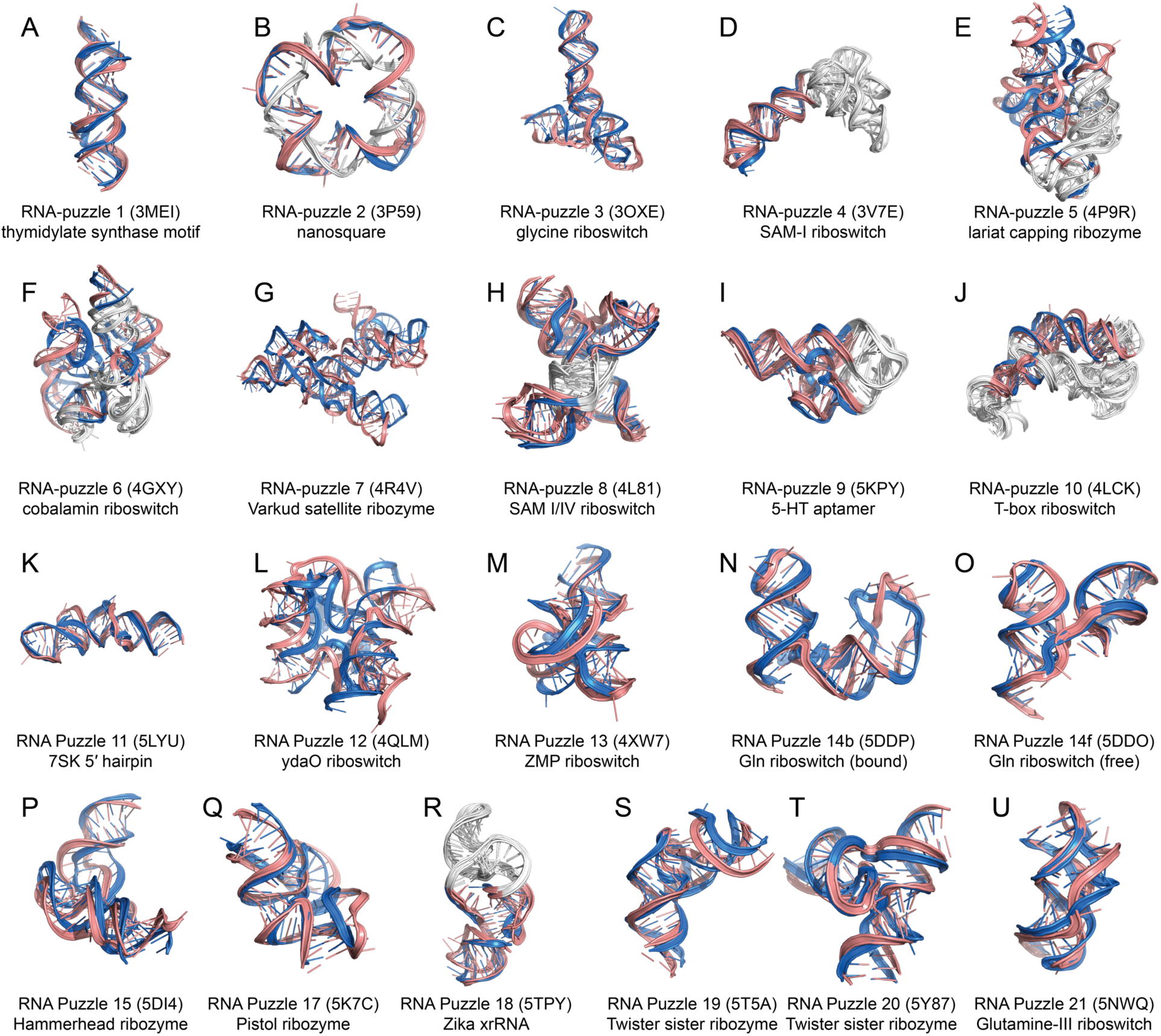
The best-of-top-1% RMSD prediction (pink) vs. native (blue) for each RNA-Puzzle in the FARFAR2-Puzzles benchmark. White regions are input template structures employed at the original time of modeling and employed in the FARFAR2 simulation.

**Table 1.** Results for each RNA-Puzzle challenge revisited in this work.

| Puzzle | PDB | Length<br>(Total)<sup>‡</sup> | Number of Structures<br>Generated | RNA | FARFAR2 |  |  |  |
| --- | --- | --- | --- | --- | --- | --- | --- | --- |
|  |  |  |  |  | Top 1%<br>Best<br>RMSD<sup>†</sup> | Best of 10 Low-E Cluster<br>RMSD<sup>†</sup> | Best RNA-puzzle<br>RMSD (All<br>Submissions)<sup>†</sup> | Best RNA-puzzle<br>RMSD (Das<br>Submissions)<sup>†</sup> |
| 1 | 3mei | 46 (46) | 14052 | thymidylate synthase motif | 2.03 | 2.50 | 3.40 | 3.40 |
| 2 | 3p59 | 60 (100) | 43431 | nanosquare | 2.28 | 2.71 | 2.30 | 2.45 |
| 3 | 3oxe | 84 (84) | 33442 | glycine riboswitch | 7.05 | 12.41 | 7.60 | 12.10 |
| 4 | 3v7e | 40 (126) | 5768 | SAM-I riboswitch | 2.43 | 2.52 | 3.40 | 4.50 |
| 5 | 4p9r | 130 (188) | 24903 | lariat capping ribozyme | 9.57 | 13.94 | 9.58 | 9.58 |
| 6 | 4gxy | 98 (158) | 28859 | cobalamin riboswitch | 9.98 | 13.08 | 12.28 | 12.28 |
| 7 | 4r4v | 185 (185) | 7963 | VS ribozyme | 15.21 | 18.52 | 20.72 | 20.72 |
| 8 | 4l81 | 71 (96) | 33086 | SAM I/IV | 4.65 | 5.23 | 4.80 | 4.80 |
| 9 | 5kpy | 40 (71) | 18660 | 5-HT aptamer | 4.54 | 4.56 | 5.86 | 7.87 |
| 10 | 4lck | 83 (177) | 5873 | T-box riboswitch | 6.31 | 6.31 | 6.78 | 6.78 |
| 11 | 5lyu | 57 (57) | 41952 | 7SK 5' hairpin | 4.43 | 6.04 | 5.22 | 7.50 |
| 12 | 4qlm | 117 (117) | 35506 | ydaO riboswitch | 11.73 | 13.32 | 10.15 | 12.63 |
| 13 | 4xw7 | 60 (60) | 20297 | ZMP riboswitch | 5.47 | 7.13 | 5.41 | 5.41 |
| 14b | 5ddp | 61 (61) | 24531 | Gln riboswitch (bound) | 5.81 | 6.88 | 5.79 | 6.59 |
| 14f | 5ddo | 61 (61) | 15112 | Gln riboswitch (free) | 3.26 | 11.85 | 6.05 | 6.05 |
| 15* | 5di4 | 68 (68) | 6123 | Hammerhead ribozyme | 4.44 | 5.98 | 5.30 | 9.83 |
| 17 | 5k7c | 58 (58) | 16529 | Pistol ribozyme | 5.03 | 6.69 | 7.13 | 7.13 |
| 18 | 5tpy | 34 (71) | 17091 | Zika xrRNA | 4.29 | 5.02 | 3.15 | 3.15 |
| 19 | 5t5a | 56 (62) | 4499 | Twister sister ribozyme | 4.86 | 5.16 | 5.50 | 10.20 |
| 20 | 5y87 | 62 (68) | 1547 | Twister sister ribozyme | 3.03 | 4.03 | 6.80 | 6.80 |
| 21 | 5nwq | 41 (41) | 48146 | Guanidinium-III riboswitch | 4.40 | 6.04 | 3.93 | 3.93 |

\*The native structure in 15 was corrupted by a crystal contact that caused a strand-swapped pairing intractable to prediction. We reconstructed the strand-unswapped monomer and rescored original submissions to present a more realistic modeling challenge.
<sup>†</sup>Heavyatom RMSD is calculated over all residues, following superposition over all residues.
<sup>‡</sup>Length is number of residues remodeled (not found in input templates); total counts template residues.

In 5 of 21 modeling challenges, FARFAR2 did not sample among its top 1% by energy a more accurate model than the original RNA-Puzzles submissions. RNA-Puzzles 12, 13, 14 (bound), and 21 represent structures of riboswitch aptamers with their small molecule ligands (cyclic diAMP, ZMP, glutamine, and guanidinium, respectively). For each of these cases, the original modeling required extensive manual curation as well as experimentally derived constraints or explicit modeling of the ligand binding site (Miao et al., 2017). Finally, the best Das lab model for RNA-Puzzle 18, the Zika xrRNA, was solved with the stepwise Monte Carlo method (Watkins et al., 2018); FARFAR2 does outperform the best original Das lab FARFAR model. These results indicate that FARFAR2 successfully automates the many *ad hoc* steps used in prior RNA-Puzzles challenges but could be improved further if ligand binding hypotheses and stepwise Monte Carlo could be incorporated into the modeling.

To compare FARFAR2 results to the original RNA-Puzzle submission process more directly, we needed some method of model selection to obtain a final set of ten from the ensemble of sampled models. Here, we reproduced the prior protocol for analyzing problems of this size within Rosetta, clustering the top 400 models with a 5.0 Å cluster radius. (Alternative clustering methods gave similar or slightly worse results; see Methods.) The best of ten clustered FARFAR2 models outperformed the original Das lab submission in 10 of 21 cases, and, despite no manual intervention in model selection, was less than 1.0 Å worse in RMSD in an additional 4. Furthermore, these selected models were native-like (RMSD accuracy better than 9.1 Å for short problems or 13.8 Å for long problems) in 16 of 21 cases.

In an actual blind modeling scenario, modelers may want to know how accurate their ensemble of FARFAR2 models is likely to be. Recently, extensions of FARFAR to building coordinates into RNA-protein (Kappel et al., 2018) and RNA-only electron density maps (Kappel et al., 2019) have suggested a promising approach. In those settings, the average pairwise RMSD between the ten lowest energy models was highly predictive of the average RMSD to native of those same ten models. We tested whether a similar relationship would apply in this setting – in the absence of electron density, where models are significantly more diverse. As shown in Figure 5, the average pairwise RMSD of the top 10 FARFAR2 cluster centers does correlate well with the average RMSD to native of those models (R^2^ of 0.84). The correlation is weaker than that observed in density guided modeling (grey points, Figure 5, R^2^ of 0.94), and the trend is shifted higher so that the same inter-model RMSD corresponds to a worse average RMSD-to-native in the density-free FARFAR2 cases. Nevertheless, the error in the above estimate is itself predictable; the standard deviation of the pairwise RMSDs among top-10 models predicts most of the variance in the RMSD to native among those models (with electron density, R^2^ = 0.90; without, R^2^ = 0.64). These relationships suggested that we would be able to predict ranges of model accuracy in real, blind prediction scenarios, a prospect that we tested in our final study.

**Figure 5.**
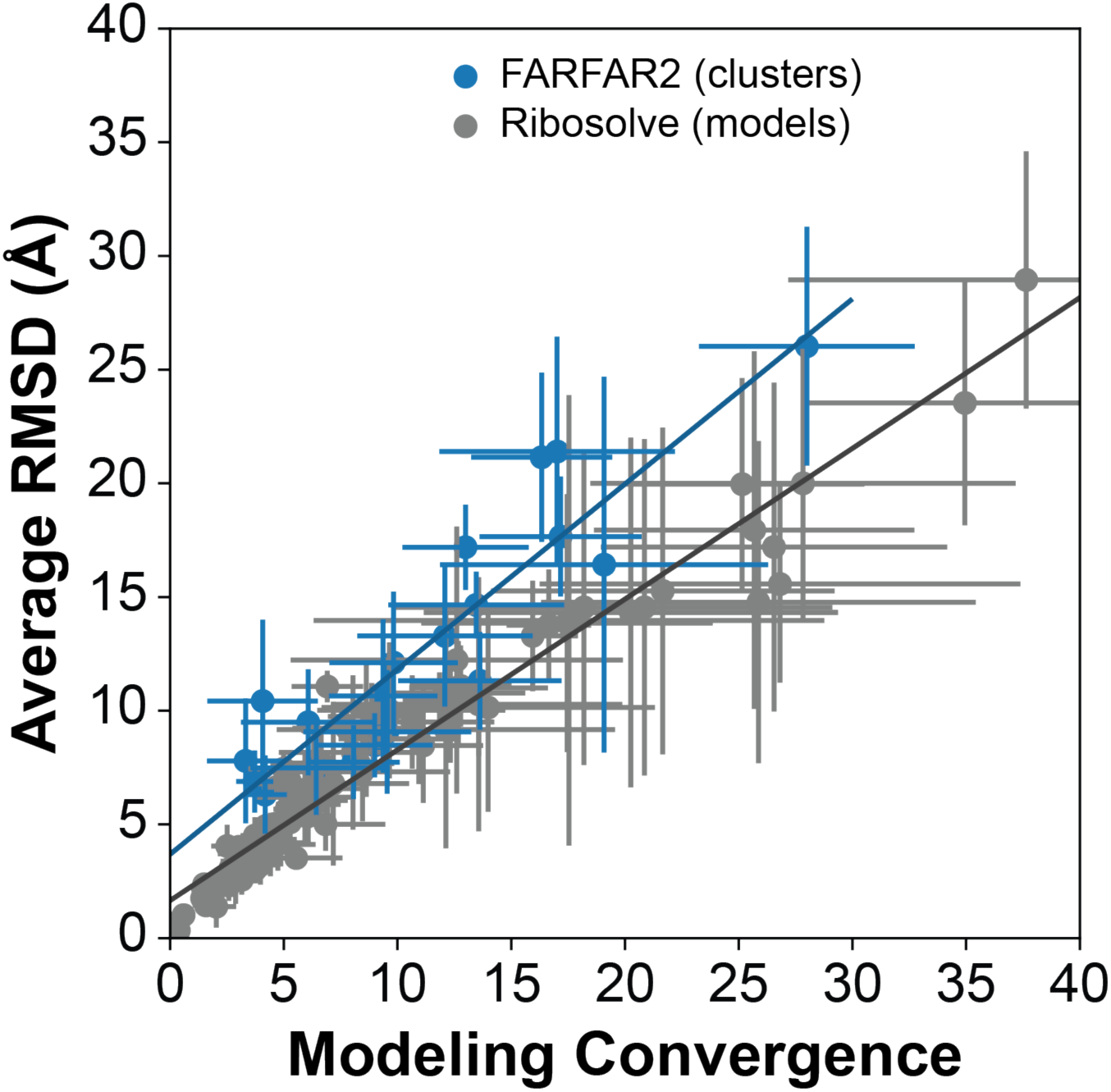
Convergence (the average pairwise RMSD among the top 10 models) is predictive of modeling accuracy (the average RMSD to native of the 10 lowest energy models or clusters) whether with electron density (DRRAFTER and Ribosolve models, gray) or without (FARFAR2, blue). Lines of best fit: Ribosolve y = 0.66 x + 1.65 Å (R^2^ = 0.94); FARFAR2 y = 0.81 x + 3.69 Å (R^2^ = 0.84). Error bars represent standard deviation of pairwise RMSD (x-error) and standard deviation of RMSD to native (y-error), which are themselves related as Ribosolve y-error = 0.76 x-error + 0.10 Å (R^2^ = 0.90); FARFAR2 y-error = 0.91 x-error + 0.09 Å (R^2^ = 0.64).

### New blind predictions of five RNA structures

After confirming modeling accuracy of FARFAR2 in the retrospective benchmarks above, we sought to validate that the FARFAR2 method was similarly effective in truly blind modeling challenges. In separate work, we have developed a pipeline for highly accurate, rapid solution of complex RNA folds using a battery of cryo-EM, multidimensional chemical mapping, and automated computational modeling (Kappel et al., 2018, 2019). We saw this as a valuable opportunity to conduct a battery of blind challenges of the FARFAR2 method. We applied FARFAR2 to predict the structures of four natural RNAs – two tandem glycine riboswitches, the S-adenosyl methionine binding SAM-IV riboswitch, and an adenoviral noncoding RNA virus-associated (VA) RNA I (Hood et al., 2019) – and a *de novo* designed RNA, a Spinach binding aptamer (Laing and Schlick, 2011). These modeling challenges were carried out fully blindly of experimental efforts by author A.M.W.; resulting models were pre-registered with the Open Science Framework (OSF) in the case of the Spinach aptamer and two glycine riboswitches; submitted to an “Unknown RFam” RNA-Puzzles challenge in the case of SAM-IV; and submitted to RNA-Puzzle 24 for VA RNA I.

The results of these five blind challenges supported the accuracy of FARFAR2 in a wide range of template-based modeling and fully *de novo* modeling scenarios. The three natural riboswitch aptamers tested use of FARFAR2 in problems where templates were available but peripheral tertiary domains needed to be built *de novo*. We used a 122 nt template derived from a crystal structure (PDB ID: 3P49) to build a series of models of the full-length 167 nt *F. nucleatum* riboswitch (Figure 5A). This template structure previously included a U1 binding loop to facilitate crystallization, which we replaced with the native sequence; it had also omitted a predicted 9-nt kink-turn linker between its two glycine-sensing domains, requiring the deletion and FARFAR2 remodeling of the first eighteen nt on the 5’ end, nt 72-91, and the three final 3’ nt, 165-167. We subsequently used part of the lowest-energy *F. nucleatum* solution, threaded with the related sequence from *V. cholerae*, to predict the structure of the 229 nt *V. cholerae* tandem glycine riboswitch, using FARFAR2 to insert a P4 stem into a multiloop of the 5’-aptamer domain and substantially remodel the resulting fold (Figure 5B). Our strategy for the SAM-IV riboswitch used a template 23-nt S-adenosyl methionine binding site from the SAM-I riboswitch (PDB ID: 2YGH) and largely followed our previously described procedure for homology modeling of complex RNA folds (Watkins et al., 2019), but used the FARFAR2 protocol instead, thus using base pair step sampling and the new fragment library in addition to the updated energy function for full-atom refinement (Figure 5C). We modeled the Spinach stabilizing aptamer Spinach-TTR-3 by employing as templates the pre-formed ideal tetraloop/receptor interaction from a P4-P6 RNA crystal structure (PDB ID: 1GID) and the DFHBI/Spinach binding site (PDB ID: 4TS2; Figure 6D). Finally, for the VA RNA I, no template structures were available, so we used only the literature secondary structure (Dzananovic et al., 2017). Upon unblinding, the modeling clearly recapitulated the global folds of the RNAs solved independently with experimental cryo-EM and mutate-and-map data (Figure 6). Using the same model selection procedure as for FARFAR2-Puzzles, we chose 10 models for each problem except the SAM-IV riboswitch, for which we could submit only 5 models. The best submitted FARFAR2 models were a 3.0 Å prediction of the *F. nucleatum* glycine riboswitch (2.2 Å over residues modeled directly), a 4.3 Å prediction of the *V. cholerae* glycine riboswitch, a 3.2 Å prediction of SAM-IV, a 6.7 Å prediction of the Spinach stabilizer (which was only determined without the ligand bound), and a 7.7 Å prediction of the VA RNA I (Table 2). Especially encouragingly, the average model accuracies were similar to predicted values: based on the mean inter-model RMSDs and the trends of Figure 5, 4.8 Å, 5.9 Å, 8.8 Å, 9.3 Å, and 13.7 Å compared to 4.9 ± 0.43 Å, 7.1 ± 1.9 Å, 12.7 ± 6.0 Å, 11.1 ± 2.9 Å, and 13.3 ± 3.4 Å, respectively. The average model was somewhat more accurate than predicted in particular for SAM-IV (8.8 Å vs 12.7 ± 6.0 Å) and Spinach-TTR-3 (9.3 Å vs. 11.1 ± 2.9 Å), but still within the range expected from the variance of FARFAR2 inter-model RMSDs.

**Figure 6.**
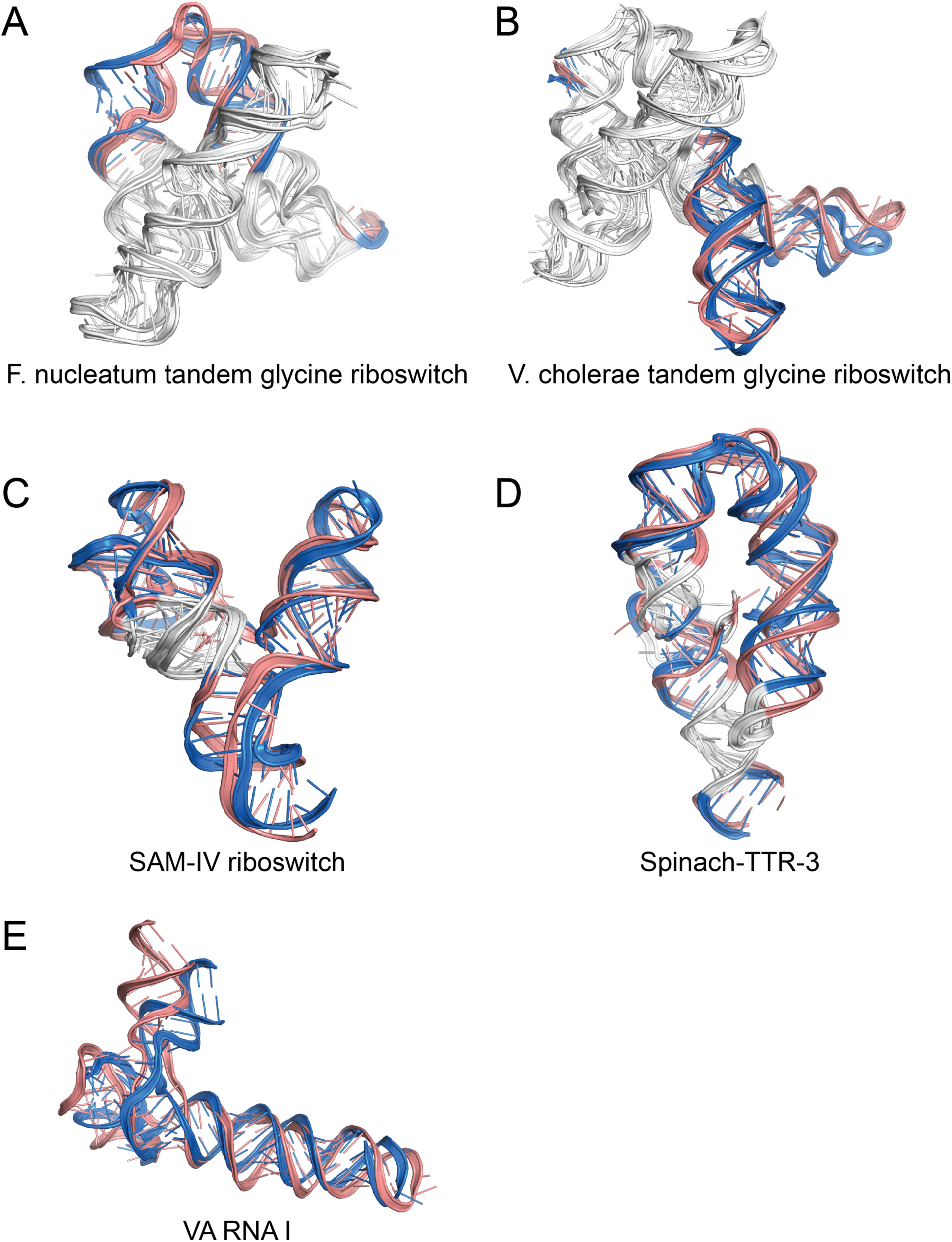
Blind predictions (salmon) of five complex RNA folds (blue) subsequently determined via cryo-electron microscopy: the (A) *F. nucleatum* and (B) *V. cholerae* full-length tandem glycine riboswitches, as well as the (C) SAM-IV riboswitch, the (D) Spinach-TTR-3, and the (E) adenoviral VA RNA I. Predictions generally achieve nucleotide accuracy. White regions are input template structures.

**Table 2.** Results for each blind challenge modeled in this work.

| Challenge | PDB | Length<br>(Total) | Number of Structures<br>Generated | RNA | Top 1%<br>Best RMSD | Best Pre-Registered<br>Model* |
| --- | --- | --- | --- | --- | --- | --- |
| 1 | pending | 45 (167) | 2681 | F. nucleatum glycine<br>riboswitch | 2.84 | 2.99 |
| 2 | pending | 145 (229) | 3949 | V. cholerae glycine<br>riboswitch | 4.42 | 4.31 |
| 3 | pending | 96 (119) | 10833 | SAM-IV riboswitch | 3.82 | 3.19 |
| 4 | pending | 97 (130) | 3336 | Spinach-TTR-3 | 6.58 | 6.66 |
| 5 | 6ol3 | 112 (112) | 87170 | VA RNA I | 4.82 | 7.66 |

\*Only five models were selected for the SAM-IV riboswitch blind challenge; ten for the others.

## Discussion

We have developed and benchmarked a major update to Rosetta fragment assembly of RNA with full-atom refinement (FARFAR2), which automates multiple steps of previous protocols and brings to bear an updated fragment library, flexible treatment of helix base pair steps, and a refined all-atom score function. We find that FARFAR2 achieves native-like structure models on three retrospective benchmarks including 18 small RNAs, 82 ‘motif-scale’ challenges, and 21 RNA-Puzzles, as well as on five new prospective blind challenges, including RNA-Puzzle 24.

Fragment assembly methods have been a classic method for structure prediction because of their simple theoretical justification: folded biomolecules occupy a special region of conformational space, and new molecules will locally resemble previously determined structures. Fragment-based methods yield reliable predictions of RNA structure, and FARFAR2’s extensions to this system – enabling kinematically realistic helix flexibility, expanding the database of crystallographic fragments, and scoring function improvements – have measurably improved its performance. Without requiring human intervention following template selection, FARFAR2 routinely samples native-like folds within its 1% lowest energy structures. These structures are consistently more accurate than the best original Das lab submissions and, in the majority of cases, more accurate than the best overall submission. Model selection based on clustering the lowest energy models appears generally successful, although our benchmark highlights cases where highly accurate models (Figure 3) are not captured by clustering. The convergence of the top ten cluster centers serves as a predictor of overall modeling accuracy. When applied to five new blind challenges, FARFAR2 achieved five blind predictions obtaining 3–8 Å accuracy on newly built residues, showing strong agreement with subsequently solved structures from cryoelectron microscopy and crystallography and confirming accuracy estimates made during modeling. FARFAR2 appears well-poised to make predictions for RNA challenges in which secondary structure and some tertiary contact information (e.g., pseudoknots) are available. Convergence-based accuracy estimates should help guide the interpretation of the models and point to cases where more experimental data are required.

The frontiers for accurate RNA 3D structure prediction involve modeling larger problems (more than 100 nucleotides of unknown tertiary structure, illustrated here by RNA-Puzzles 12, 5, and 7, Figure 4) and problems in which very high accuracy is required (e.g., if the structure is required for subsequent ligand docking or drug design). Fully automated methods that more accurately solve subproblems at the expense of more computation, such as stepwise Monte Carlo (Watkins et al., 2018), may bridge these gaps if pipelined with fragment assembly, as may methods that make more active use of residue contact inference from sequence covariation (Weinreb et al., 2016) or artificial neural networks (AlQuraishi, 2019; Li et al., 2019). Beyond the FARFAR2 algorithm itself, which enables largely automated prediction of RNA structure with good accuracy, this benchmark system and the associated ‘decoy’ models in the FARFAR2-Classics, FARFAR2-Motifs, and FARFAR2-Puzzles data sets can serve as starting points for the development of the next generation of high-resolution structure prediction algorithms.

## Acknowledgements

We thank K. Kappel, Z. Su, K. Zhang, and W. Chiu (Stanford) for independent structure determination of three new blind challenges. We thank Z. Miao and E. Westhof for organizing the RNA-puzzles and ‘Unknown RFam’ challenges. We acknowledge funding from the National Institutes of Health (R21 CA219847 and R35 GM122579) and the Army Research Office W911NF-16-1-0372.

## Author Contributions

Conceptualization, A.M.W. and R.D.; Methodology, A.M.W. and R.D.; Investigation, A.M.W.; Writing – Original Draft, A.M.W.; Writing – Review & Editing, A.M.W. and R.D.; Funding Acquisition, R.D.; Resources, R.D.; Supervision, R.D.

## STAR Methods

### CONTACT FOR REAGENT AND RESOURCE SHARING

Further information and requests for resources and reagents should be directed to and will be fulfilled by the Lead Contact, Rhiju Das.

### METHOD DETAILS

All Rosetta commands were run using Rosetta weekly release 2019.22, available for free download for noncommercial use at http://www.rosettacommons.org.

#### An automated fragment assembly benchmark

Two technical improvements within Rosetta permitted rapid progress on a fully automated fragment assembly protocol. First, FARFAR2 jobs may be fully specified and run using a single command line, rather than requiring pre-configuration using a ‘params file’ with a complex language (Das and Baker, 2007); example command-lines are given below (‘FARFAR2 execution on benchmark cases’). Second, we developed a benchmarking framework (Watkins et al., 2018), available at https://github.com/daslab/rna_benchmark with documentation and instructions.

#### New fragment set

We obtained release 3.10 of representative nonredundant 3D structures from BGSU’s RNA 3D Hub (Leontis and Zirbel, 2012). We obtained the indicated PDB chains and parsed them into fragment and jump (base-base rigid body transformations for base pairs) database files using the following command lines:

~~~
rna_database -s *_RNA.pdb -vall_torsions
RICHARDSON_RNA18_2.5_revised.torsions -cut_at_rna_chainbreak true
-ignore_zero_occupancy false -guarantee_no_DNA true
~~~

~~~
rna_database -s *_RNA.pdb -jump_database true -cut_at_rna_chainbreak true
-ignore_zero_occupancy false -guarantee_no_DNA true -out:file:o
RICHARDSON_RNA18_2.5_revised.jump
~~~

#### Fragment homology exclusion

Benchmarks of fragment assembly approaches can give misleading overestimates of accuracy if fragments of the experimental (‘native’) structure are included during modeling. In order to better mimic blind prediction scenarios in which the experimental structure is not available at the time of modeling, we ensured that our fragment libraries were free of contamination from the native structures employed in the benchmarks through a “fragment homology exclusion” option implemented in FARFAR and controlled by command-line options; see also (Watkins et al., 2018). In this mode, all six-nucleotide contiguous stretches of RNA to be built are extracted from the experimental structure, and any fragments in the fragment library that are deemed too similar to the experimental structure are excluded as possible contamination from that experimental structure or a close homolog. More specifically, these ranges of structure are compared by heavy-atom RMSD to every fragment in the fragment library with matching purine/pyrimidine content, and fragments with RMSD less than 1.2 Å from the experimental structure are eliminated from consideration.

#### Modes of applying secondary structure information

In most modeling cases we have encountered, models of secondary structure were previously available based on expert analysis, thermodynamic modeling packages (Hofacker and Lorenz, 2014; Mathews, 2006), and/or sequence alignment information. Such known secondary structure (see Supplemental Table 4) can now be specified either via command-line or in an input text file. The FARFAR2 protocol can apply this secondary structure information in four ways. First, to replicate the original FARNA style of simulation, one may specify no secondary structure at all. Second, a specified secondary structure can provide energetic “base pair constraints” (see **FARFAR2 execution on benchmark cases** below) that tend to draw paired residues together, first introduced in (Das et al., 2010). Base pair constraints consist of harmonic restraints placed on the distance between corresponding Watson-Crick edge atoms (1.9 Å with a standard deviation of 0.1 Å) as well as a harmonic restraint placed on the distance between sugar C1 atoms (10.5 Å with a standard deviation of 1 Å) to favor paired, rather than stacked-and-tilted, conformations. The influence of these constraints are, by default, ramped up over each round of low-resolution modeling, then applied at full strength during minimization. Third, helical stems may be generated and then provided as fixed inputs to the simulation (either individual helices or ensembles thereof), as described in (Cheng et al., 2015; Miao et al., 2015). This procedure is automated by the `rna_benchmark` system but may be carried out manually using scripts from the Rosetta tools directory (see **Setting up helices** below). Fourth, helix flexibility may be simulated directly in a kinematically realistic way by sampling from a library of “base pair steps”. Base pair step moves keep the base paired secondary structure fixed while sampling orientation changes between consecutive base pairs seen in the crystallographic database; Rosetta’s implementation is generalized to permit realistic sampling of stems with interrupting nucleotide bulges on one side, as well. This mode had not been previously described or tested but is now the default mode, due to its superior performance in the benchmarking described in the main text and supplement. Detailed comparison of the effects of base pair step sampling on Puzzle 21, a guanidinium riboswitch, is shown in Figure S4; a comparison of its performance versus fixed helices across multiple large puzzles is depicted in Figure S5.

#### Benchmark cases

The three benchmarks evaluated were intended to evaluate the performance of the new FARFAR2 ‘best practices’ on challenges of qualitatively different scales. The original FARNA benchmark and the stepwise Monte Carlo benchmark examined relatively small structures, though many representative examples in the latter case were fairly complex. In contrast, the RNA-Puzzles benchmark, like the five blind challenges undertaken to test this method, examined entire folded RNAs, typically with many tertiary contacts.

The FARFAR2-Classics benchmark comprised 18 challenges, each of which was either a single-stranded stem-loop or a duplex. Two structures that overlapped exactly with the FARFAR2-Motifs benchmark (see below) were omitted from the set of 20 structures (1J6S, a G-quadruplex; 1ZIH, a GCAA tetraloop) used to benchmark the original FARNA algorithm (Das and Baker, 2007). In each case, the challenge structure was the entire crystallized RNA. As the central benchmark challenges, these were approached in four distinct modes of secondary structure specification: none, “base pair constraint” generation, fixed helical “chunks,” and “base pair step” sampling. Either the default fragment library or a newly generated fragment library were used to sample loop nucleotides and, if applicable, base pair steps. Finally, results for each challenge were either left un-minimized or optimized in one of two scoring functions: the original FARFAR refinement scoring function (hereafter “hires”) or the modern RNA scoring function developed for stepwise Monte Carlo (originally termed *rna_res_level_energy4.wts*, hereafter called “res4”) (Watkins et al., 2018). (The same low-resolution structures were minimized to produce the final data set.) As in the original FARNA paper, 50,000 structures were generated for each simulation. Additionally, as a control, we replicated the exact parameters of the original FARNA work, including an eleven-year-old fragment library based only on the *H. marismortui* 23S ribosome (PDB: 1JJ2). This replication performed better on the majority of cases, suggesting some level of Rosetta simulation improvement and bug fixes since 2007 (see Figure S1, Table S1). For this comparison, we had to reduce the fragment homology exclusion radius from 1.2 Å to 0.5 Å or else we were unable to discover fragments for cases 1CSL, 1DQF, 1ESY, 1I9X, 1KD5, 1Q9A, 1QWA, 1XJR, 2A43, or 2F88. Informed by a comparison of these simulation conditions, we decided to take the SWM “res4” scoring function and the new fragment libraries as the FARFAR2 standard. We also chose to specify secondary structure through base pair steps. Though the advantage over fixed helical stems was not obvious across the FARNA benchmark, and we anticipated that more complex tertiary environments, such as those found in typical whole structure prediction cases, would benefit further from some form of helix flexibility, and this was later confirmed (see main text; and Figures S4-S5). These optimal FARFAR2 parameters also proved robust to variations in other simulation settings. We repeated the whole benchmark set with the final FARFAR2 settings but disabled all low-resolution filters (which restart simulations that have generated structures with chainbreaks, missing base pairs, or bad scores). We also tested reducing the ten-round fragment assembly schedule to only one or two rounds, and we still obtained excellent results (Supplemental Figure S1D). None of these configuration variants ought to be taken as new best practices as their own, however, particularly for larger RNA folds. Setting up only one simulation round, for example, increases the computational expense more than fourfold on average, and while score, base pair, and chainbreak filters may not be very important for simple folds that are quick to energy-minimize, they prevent considerable needless computation on large RNAs.

The FARFAR2-Motifs benchmark set was chosen because it was a direct expansion of the earlier FARFAR benchmark (Das et al., 2010), augmented with challenges that demonstrated new features of FARFAR2 (such as ‘aligned’ cases, which permit expert specification of the relative orientation of a subset of the native structure; and support for chemically modified nucleotides). This expansion had originated when developing the stepwise Monte Carlo (SWM) algorithm and included single-stranded loops, two-way and multi-way junctions, tertiary contacts, and motifs that exist outside of a Watson-Crick context (such as quadruplexes (Pan et al., 2006) and parallel strands (Safaee et al., 2013)). We found that SWM frequently produced excellent native RMSDs. We generated 10,000 structures for each benchmark case: fewer than for FARFAR2-Classics because most of the above structures were smaller and structurally simpler, and because there are more than four times as many benchmark cases. Though SWM delivers performance unattainable by fragment methods, strong performance on this benchmark set would suggest the ability of fragment assembly methods to provide good starting points for higher-resolution simulation at a fraction of the computational cost required.

The FARFAR2-Puzzles benchmark set includes all RNA-Puzzles for which solutions and submissions were available at the time of this study: that is, all of the first 21 except for 16, and including both bound and unbound states of 14. In some cases, a large proportion of the problem was known approximately by homology, while in others only the helices alone were an appropriate starting point. Each problem was run to generate several thousand structures, aiming for substantial sampling at modest computational expense. (Since typical RNA-Puzzle rounds allow three to four weeks of prediction, this is a fraction of the total sampling available for a typical challenge.)

#### FARFAR2 execution on benchmark cases

We tested multiple combinations of settings for the FARFAR2 protocol. We present here the relevant command lines for each possible configuration, as operated on the 157D challenge from the FARNA benchmark. (The command line argument to run with a particular set of conditions, such as secondary structure specification vs. number of rounds, are mutually independent of each other and can be recombined at will.)

##### New recommended defaults

Before delving into the details of benchmark conditions, here is an example of how one might specify a modeling problem using the new FARFAR2 defaults. Suppose one is concerned with the guanidinium-III riboswitch. First, prepare a FASTA-formatted file `guanidine_III.fasta`:

~~~
>guanidine_III A:1-41
ccggacgaggugcgccguacccggucaggacaagacggcgc
~~~

Then, prepare a file containing the dot-bracket notation secondary structure, `guanidine_III.secstruct`. Since only the first line of the secondary structure file is read, one may include the sequence as the second line as a useful point of reference.

~~~
[[[[.......(((((((..]]]]..........)))))))
ccggacgaggugcgccguacccggucaggacaagacggcgc
~~~

Finally, execute the following command:

~~~
rna_denovo -fasta guanidine_III.fasta -secstruct_file guanidine_III.fasta
-minimize_rna true
~~~

One may provide a parameter `-nstruct` to control how many structures are generated by each execution of the above command; the parameter `-out:file:silent` names the output file where results are accumulated.

##### FARNA replication

An exact replication of the original FARNA benchmark required a special flag to use the original fragment library using only a single ribosome crystal structure (PDB 1JJ2). The specification of a single base pair (as found in `157d_orig_START1_157d.pdb`) is the minimum necessary information to seed simulation of a bimolecular RNA.

~~~
rna_denovo -fasta 157d_orig.fasta -native 157d_orig_NATIVE_157d.pdb
-s 157d_orig_START1_157d.pdb -minimize_rna false -cycles 20000 -nstruct 20
-use_1jj2_torsions true
~~~

##### No secondary structure specification, no minimization, old fragments

This run replicates the FARNA benchmark conditions but uses the fragment set that was in regular use by 2010.

~~~
rna_denovo -fasta 157d_orig.fasta -native 157d_orig_NATIVE_157d.pdb
-s 157d_orig_START1_157d.pdb -minimize_rna false -cycles 20000 -nstruct 20
-fragment_homology_rmsd 1.2 -exclusion_match_type MATCH_YR
-jump_library_file 1jj2_RNA_jump_library.dat
-vall_torsions RICHARDSON_RNA09.torsions -bps_moves false
~~~

##### No secondary structure specification, no minimization, new fragments

This run uses a new set of fragments generated for this work.

~~~
rna_denovo -fasta 157d_orig.fasta -native 157d_orig_NATIVE_157d.pdb
-s 157d_orig_START1_157d.pdb -minimize_rna false -cycles 20000 -nstruct 20
-fragment_homology_rmsd 1.2 -exclusion_match_type MATCH_YR -bps_moves false
~~~

##### No secondary structure specification, minimization under ‘rna_hires.wts’, new fragments

This run uses the new set of fragments generated for this work.

~~~
rna_denovo -fasta 157d_orig.fasta -native 157d_orig_NATIVE_157d.pdb
-s 157d_orig_START1_157d.pdb -minimize_rna true -cycles 20000 -nstruct 20
-fragment_homology_rmsd 1.2 -exclusion_match_type MATCH_YR
-score:weights rna/denovo/rna_hires.wts -bps_moves false
~~~

##### No secondary structure specification, minimization under ‘rna_res_level_energy4.wts’, new fragments

This run uses a new set of fragments generated for this work.

~~~
rna_denovo -fasta 157d_orig.fasta -native 157d_orig_NATIVE_157d.pdb
-s 157d_orig_START1_157d.pdb -minimize_rna true -cycles 20000 -nstruct 20
-fragment_homology_rmsd 1.2 -exclusion_match_type MATCH_YR -bps_moves false
~~~

##### Base pair constraints, minimization under ‘rna_res_level_energy4.wts’, new fragments

This run specifies a secondary structure to generate base pair constraints; it does not require a starting PDB.

~~~
rna_denovo -fasta 157d_orig.fasta -native 157d_orig_NATIVE_157d.pdb
-secstruct “(((.((((.(((,))).)))).)))” -minimize_rna true -cycles 20000
-nstruct 20 -fragment_homology_rmsd 1.2 -exclusion_match_type MATCH_YR - bps_moves false
~~~

##### Base pair steps, minimization under ‘rna_res_level_energy4.wts’, new fragments

This run specifies a secondary structure to sample base pair steps; it does not require a starting PDB.

~~~
rna_denovo -fasta 157d_orig.fasta -native 157d_orig_NATIVE_157d.pdb
-secstruct “(((.((((.(((,))).)))).)))” -minimize_rna true -cycles 20000
-nstruct 20 -fragment_homology_rmsd 1.2 -exclusion_match_type MATCH_YR
~~~

##### Fixed helical inputs, minimization under ‘rna_res_level_energy4.wts’, new fragments

This run uses multiple fixed input stems.

~~~
rna_denovo -fasta 157d_orig.fasta -native 157d_orig_NATIVE_157d.pdb
-s 157d_chunks_HELIX1_157d.pdb 157d_chunks_HELIX2_157d.pdb
157d_chunks_HELIX3_157d.pdb -minimize_rna true -cycles 20000 -nstruct 20
-fragment_homology_rmsd 1.2 -exclusion_match_type MATCH_YR -bps_moves false
~~~

##### Base pair steps, minimization under ‘rna_res_level_energy4.wts’, new fragments, no filters

This run turns off filters (for score and chain closure) used during the low resolution phase to improve the quality of output structures at the cost of additional run time.

~~~
rna_denovo -fasta 157d_orig.fasta -native 157d_orig_NATIVE_157d.pdb
-secstruct “(((.((((.(((,))).)))).)))” -minimize_rna true -cycles 20000
-nstruct 20 -fragment_homology_rmsd 1.2 -exclusion_match_type MATCH_YR
-no_filters
~~~

##### Base pair steps, minimization under ‘rna_res_level_energy4.wts’, new fragments, one round of fragment assembly

Typical runs of fragment assembly go through ten rounds: three rounds of 3-mer fragment insertion, three of 2-mers, and four of 1-mers, while ramping the weights of particular score terms. Setting the number of total rounds to one or two makes the protocol much more abrupt in its transitions but does not affect other simulation settings.

~~~
rna_denovo -fasta 157d_orig.fasta -native 157d_orig_NATIVE_157d.pdb
-secstruct “(((.((((.(((,))).)))).)))” -minimize_rna true -cycles 20000
-nstruct 20 -fragment_homology_rmsd 1.2 -exclusion_match_type MATCH_YR
-rounds 1
~~~

#### Setting up helices

To use fixed helical inputs for a FARFAR structure prediction challenge without using the rna_benchmark setup scripts, a Python script is available within Rosetta tools:

~~~
rna_helix.py -seq ggg ccc
~~~

will create a three base pair helix of sequence GGGCCC, numbered 1-6 with no chain letter. To impose some reasonable numbering:

~~~
renumber_pdb_in_place.py helix.pdb A:1-3 B:1-3
~~~

#### Sources of experimental PDB structures

The structure sources for the motif benchmark have been described previously. (Watkins et al., 2018) The structures for the FARNA benchmark are as follows:

PDB codes 157D (Leonard et al., 1994); 1A4D (Dallas and Moore, 1997); 1CSL (Ippolito and Steitz, 2000); 1DQF (Xiong and Sundaralingam, 2000); 1ESY (Amarasinghe et al., 2000); 1I9X (Berglund et al., 2001); 1KD5 (Kacer et al., 2003); 1KKA (Cabello-Villegas et al., 2002); 1L2X (Egli et al., 2002); 1MHK (Szép et al., 2003); 1Q9A (Correll et al., 2003); 1QWA (Finger et al., 2003), 1XJR (Robertson et al., 2005); 255D (Holbrook et al., 1991); 283D (Baeyens et al., 1996); 28SP (Schmitz et al., 1999); 2A43 (Pallan et al., 2005), 2F88 (Seetharaman et al., 2006).

The structures for the RNA-Puzzles benchmark are:

PDB codes 3MEI; 3MEI (Dibrov et al., 2011a); 3P59 (Dibrov et al., 2011b); 3OXE (Huang et al., 2010); 3V7E (Baird et al., 2012); 4P9R (Meyer et al., 2014); 4GXY (Peselis and Serganov, 2012); 4R4V (Suslov et al., 2015); 4L81 (Trausch et al., 2014); 5KPY (Porter et al., 2017); 4LCK (Zhang and Ferré-D’Amaré, 2013); 5LYU (Martinez-Zapien et al., 2017); 4QLM (Ren and Patel, 2014); 4XW7 (Trausch et al., 2015); 5DDP (Ren et al., 2015); 5DDO (Ren et al., 2015); 5DI4 (Mir et al., 2015); 5K7C (Ren et al., 2016); 5TPY (Akiyama et al., 2016); 5T5A (Liu et al., 2017); 5Y87 (Zheng et al., 2017), 5NWQ (Huang et al., 2017).

Structures used as templates in the blind predictions: 3P49 (Butler et al., 2011), 2YGH (Schroeder et al., 2011), 4TS2 (Warner et al., 2014), 1GID (Cate et al., 1996).

### QUANTIFICATION AND STATISTICAL ANALYSIS

#### Data analysis

FARFAR2 simulations produce a compressed Rosetta-format file called a ‘silent file’ representing each trajectory endpoint. These files may be turned into PDB-format coordinate files using a Rosetta executable, `extract_pdbs`. They also hold scoring information; each line beginning with `SCORE:` either describes what scoring terms were used in generating the silent file or the values for that particular structure. Using these ‘score-lines’, the programs `grep` and `awk`, and the GNU coreutils `sort` and `wc`, we sorted the silent file by total score and by RMSD, and thereby selected the best-RMSD structure, the best-score structure, and the best-RMSD structure from among the top 1% by RMSD. As an example, the following command sorts the ‘score-lines’ by total energy, takes the top 500 lines, re-sorts those lines by RMSD, and prints out the ‘tag’ of the best-RMSD structure from those top 500 by score.

~~~
grep “^SCORE:” farna_rebuild.out | grep -v description | sort -nk2 | head -n
500 | sort -nk24 | head -n 1 | awk ‘{print $NF}’
~~~

The following Rosetta command will extract that specified model `$TAG` as a PDB file called `$TAG.pdb`.

~~~
extract_pdbs -in:file:silent farna_rebuild.out -tags $TAG
~~~

#### Model selection

In order to produce a selection of models for comparison to RNA-Puzzles submissions, we attempt a variety of clustering strategies. The following command line will cluster the lowest energy `$NN` models with a `$RR` Å radius:

~~~
rna_cluster -in:file:silent
farna_rebuild.out -out:file:silent farna_rebuild.clustered.out -native 157d_orig_NATIVE_157d.pdb -nstruct $NN
-cluster_radius $RR
~~~

We chose three values for `$NN`: 400 (the program default), 1% of the decoys, or 2% of the decoys. We also surveyed possible values for `$RR`: for constant values, we chose 2.0, 3.0, 5.0, and 7.0; we also employed an ‘adaptive’ clustering radius that responds to how tightly converged the top models are for a trajectory – for this radius, we chose 0.5, 1.0, or 1.5 times the average pairwise RMSD among the top ten models for the run in question. A 5.0 Å radius and 400 models were the original choice and also ended up being the best-performing choice by a slight margin.

For comparisons with the original FARNA algorithm analysis of the FARFAR2-Classics benchmark, the above clustering command was used with `$NN` = 5000 and `$RR` = 3.0 (Das and Baker, 2007). For comparisons with the original SWM execution of the FARFAR2-Motifs benchmark, the above clustering command was used with `$NN` = 400 and `$RR` = 2.0 (Watkins et al., 2018).

#### Measures of prediction accuracy

We measure prediction accuracy in the benchmarks using heavy-atom root-mean-squared deviation (RMSD). RMSD is obtained by aligning the model structure to the native structure and taking the average distance between all heavy-atoms (i.e., non-hydrogens) in the structure.

#### Translating RMSD thresholds

The RMSD_100_ measure allows the comparison of RMSDs on different structure sizes by scaling to 100 residues (Carugo and Pongor, 2008):

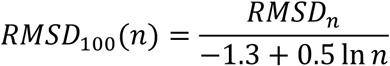

An extension of this measure (Kappel and Das, 2019) permits scaling between any numbers of residues:

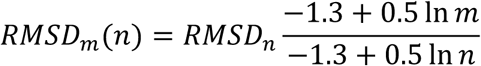

We applied this to convert RMSD 4.0 Å (the threshold treated as ‘native-like’ in the original FARNA publication and used for the *FARFAR2-Classics* benchmark in this work) to a threshold for *FARFAR2-Puzzles*. The *FARFAR2-Classics* benchmark had a median length of 26 moving residues; the median *FARFAR2-Puzzles* case featured 71. Because of the presence of a few long RNA-Puzzle problems, we also separately considered 18 “short problems” with median 68 moving residues, and 3 “long problems” with median 130 moving residues. Applying the above formula to convert (4.0 Å, 26) yields (9.2 Å, 71) or else (9.1 Å, 68); (13.8 Å, 130). We use the tighter 9.1 Å threshold for either “overall” or short problems alone; and 13.8 Å for the largest three problems. These thresholds also corresponded well to visual assessment of whether models were ‘native-like’ for each of the size ranges (see, e.g., Figures 2, 4, and 5).

### DATA AND SOFTWARE AVAILABILITY

Applications for running FARFAR2 and scripts for working with PDB files for use with this algorithm are part of the Rosetta software suite, which is free for academic use (https://rosettacommons.org). FARFAR2 is available as a webserver on ROSIE at https://rosie.rosettacommons.org/farfar2. The scripts used to set up the benchmarks studied in this work are available at https://github.com/DasLab/rna_benchmark. The final raw dataset is deposited with Stanford Library’s PURL system at https://purl.stanford.edu/wn364wz7925.

### KEY RESOURCES TABLE

The key resources table has been provided as a separate attachment.

