## Supplemental Figures and Tables for "FARFAR2: Improved de novo Rosetta prediction of complex global RNA folds"

### Supplemental Information for "FARFAR2: Improved de novo Rosetta prediction of complex global RNA folds"

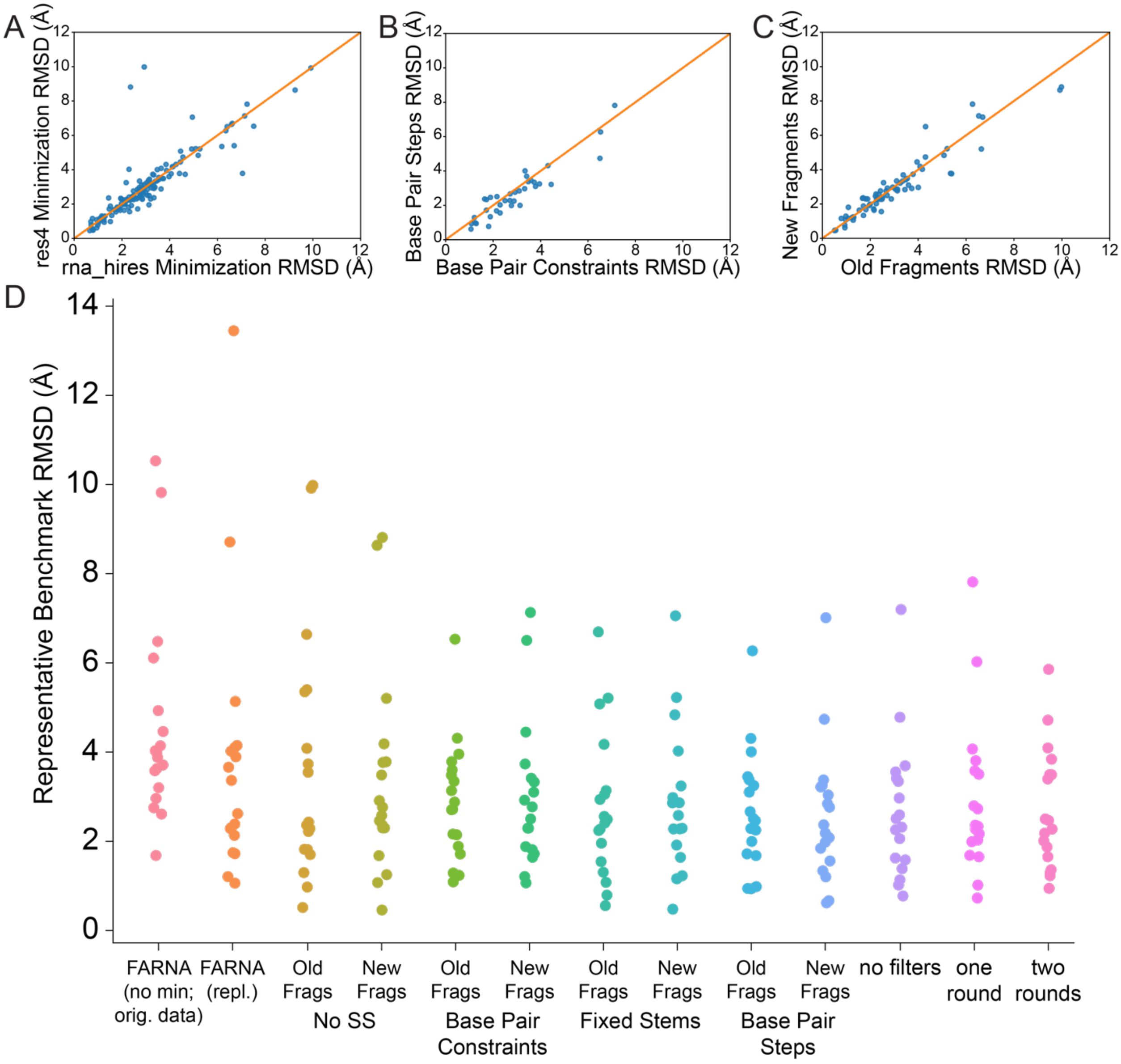

**Figure S1.** Related to Figure 1. FARFAR2 performance on the Classics benchmark. (A) Optimization in the energy function originally developed for SWM was more effective than the original FARFAR energy function (better RMSD in 91 of 144 possible comparisons) and considerably more effective than no minimization at all (better RMSD in 102 cases, often by considerable margins). (B) Providing secondary structure information, particularly as fixed helices or base pair steps, offered substantial advantages over energetic restraints or no secondary structure information. We compared top 1% RMSD values, holding other simulation conditions constant, and only considering the SWM energy function for brevity; fixed stems produced a superior RMSD in 68 of 108 possible such comparisons against other secondary structure specification methods, while base pair steps produced a superior RMSD in 76 of 108 comparisons. (C) Some benefit was also seen by using a newly obtained fragment library (43 of 72 comparisons; greatest advantage seen in hardest problems). (D) A full comparison of simulation results using “res4” minimization, versus FARNA simulation and controls using the FARFAR2 simulation parameters but no filters or only one or two rounds of fragment assembly.

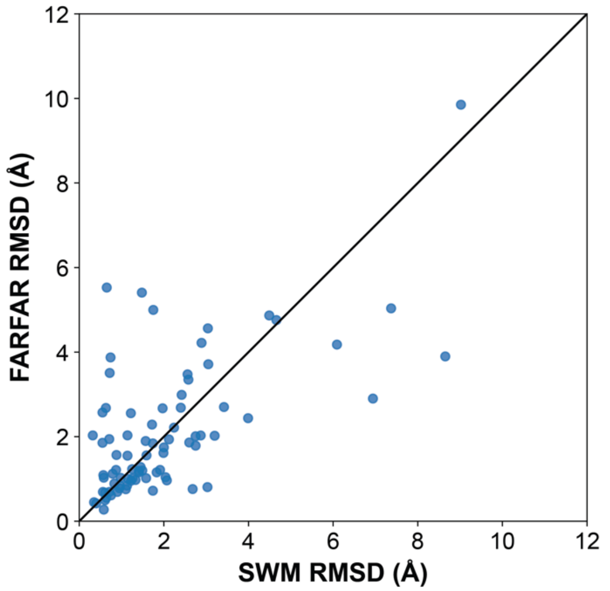

**Figure S2. Related to Figure 1.**

FARFAR2 yields a superior RMSD to SWM among its five low-energy cluster centers in 44 of 82 benchmark cases. FARFAR2's particular advantage lies in cases where *both* SWM and FARFAR2 fail to obtain 1.5 Å RMSD accuracy: of the 6 cases where SWM provides worse than 5.0 Å RMSD, FARFAR2 provides a superior RMSD in 5. In contrast, among the 42 cases where SWM achieves better than a 1.5 Å RMSD, FARFAR2 obtains superior RMSD in only 19.

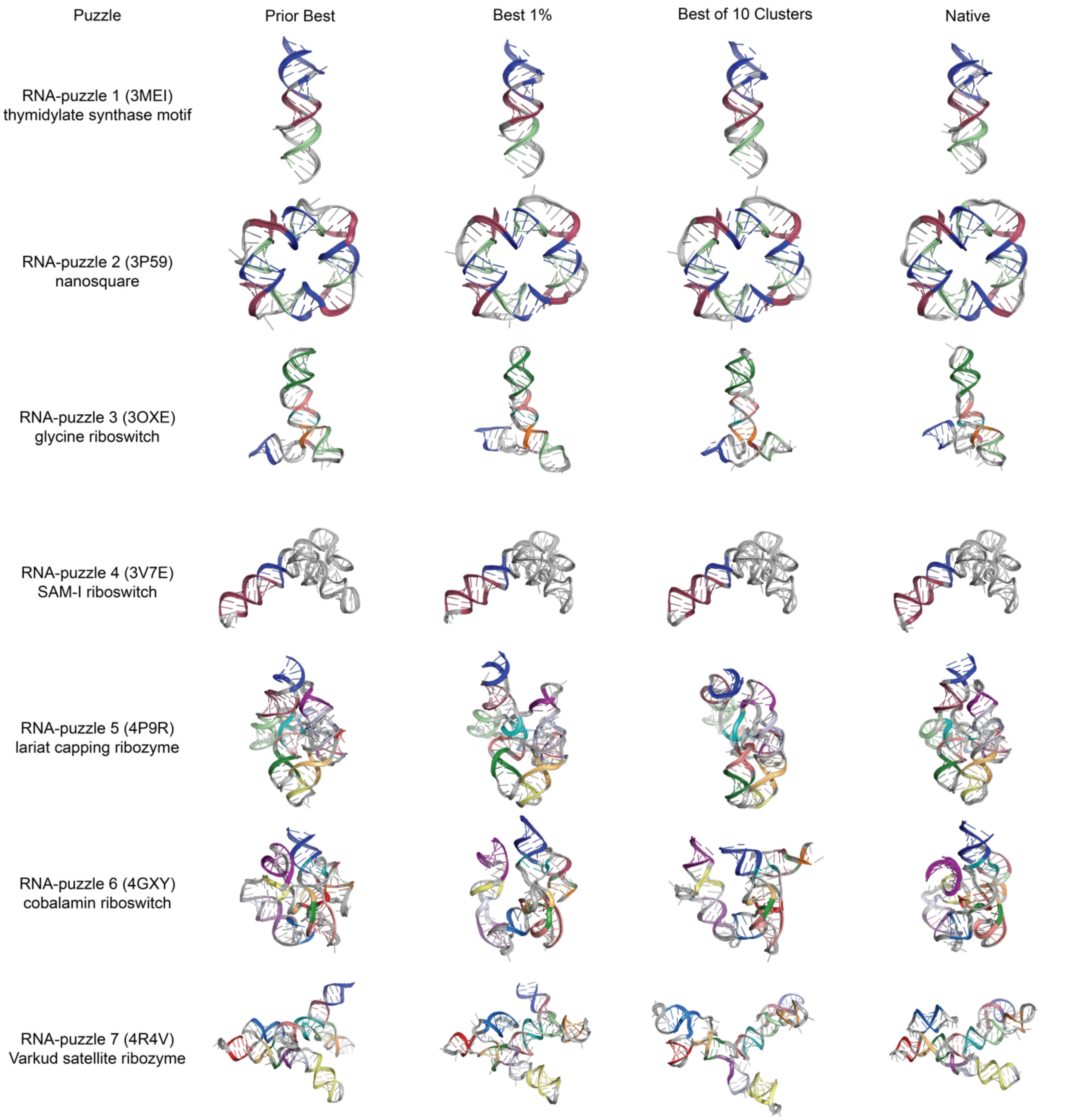

**Figure S3.** Related to Figure 4. Detailed depictions of each FARFAR2-Puzzles benchmark case, including the best originally submitted model, the FARFAR2 model with lowest RMSD in the top 1% of models overall, the lowest RMSD cluster center among the top 10 by energy, and the native structure. Models are colored to highlight distinct secondary structure elements. (1 of 3)

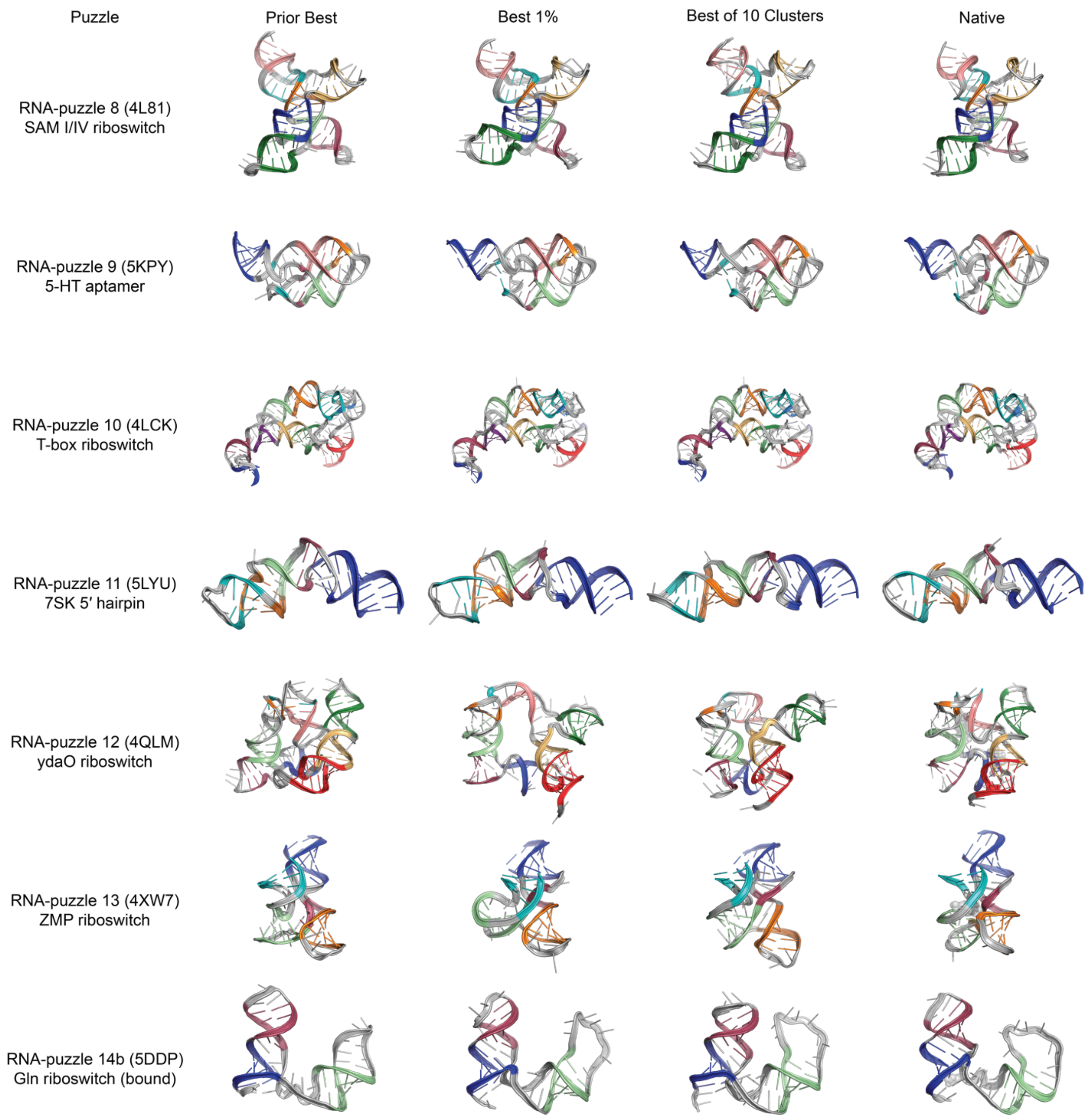

**Figure S3.** Related to Figure 4. Detailed depictions of each FARFAR2-Puzzles benchmark case, including the best originally submitted model, the FARFAR2 model with lowest RMSD in the top 1% of models overall, the lowest RMSD cluster center among the top 10 by energy, and the native structure. Models are colored to highlight distinct secondary structure elements. (2 of 3)

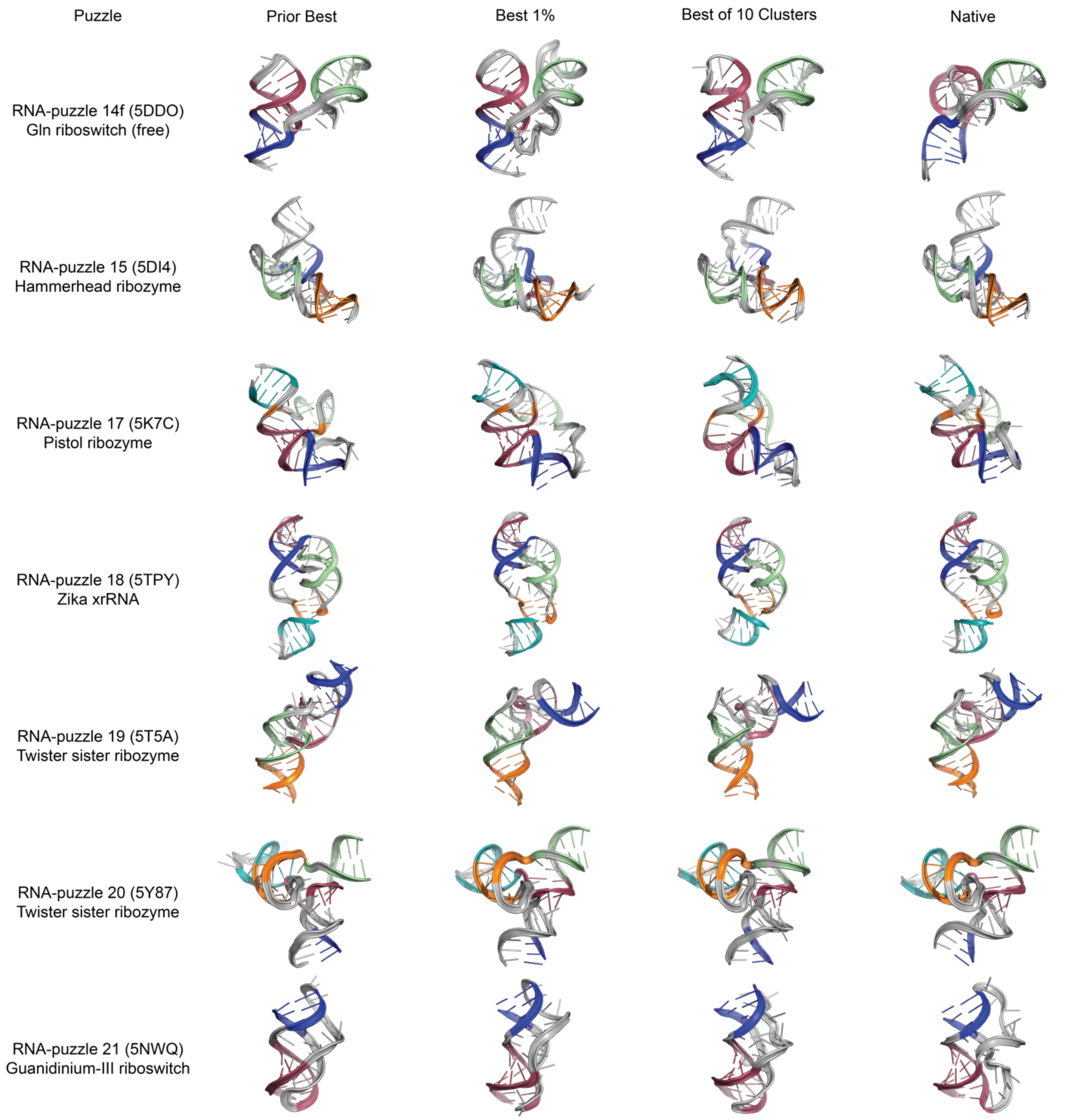

**Figure S3.** Related to Figure 4. Detailed depictions of each FARFAR2-Puzzles benchmark case, including the best originally submitted model, the FARFAR2 model with lowest RMSD in the top 1% of models overall, the lowest RMSD cluster center among the top 10 by energy, and the native structure. Models are colored to highlight distinct secondary structure elements. (3 of 3)

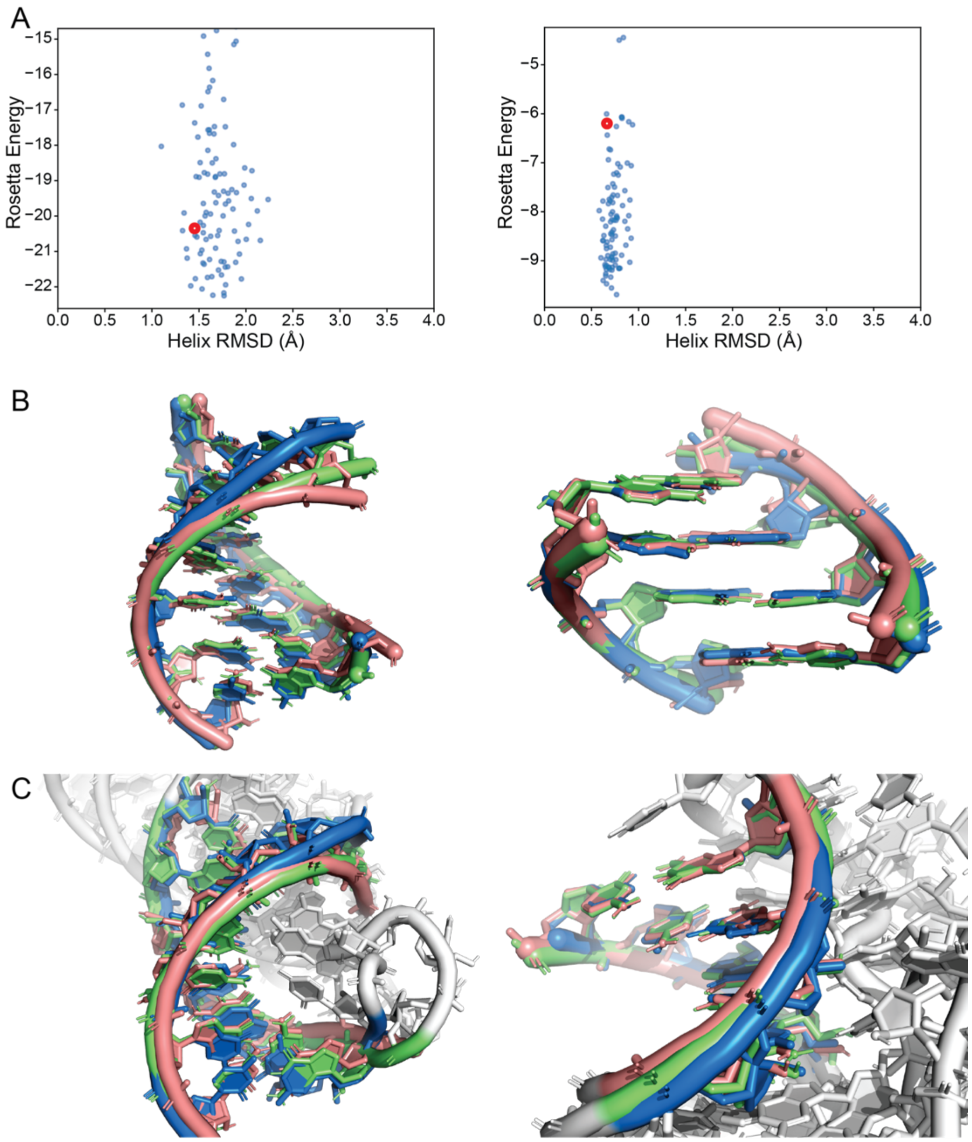

**Figure S4.** Related to Figure 3. Base pair steps provide the benefits of pre-generated helical ensembles. (A) Even without surrounding RNA context to help guide helix flexibility toward the native conformation, the base pair step helices (blue) obtained superior native RMSD and energy to the fixed stems (red) in both cases. (B, C) RNA-Puzzle 21 (blue) was also approached using fixed stems (pink) versus base pair step sampling (green), and for both helices, fixed stems yielded a worse RMSD to the native helix conformation (1.6 Å > 1.3 Å; 0.8 Å > 0.5 Å). These improvements represented geometrically important flexibility: the tightly pseudoknotted structure of Puzzle 21 features no external stacking on either helix, and deviations from ideality are essential within each helix in order to relieve strain and achieve conformations where the helix termini have no residues stacked upon them (C).

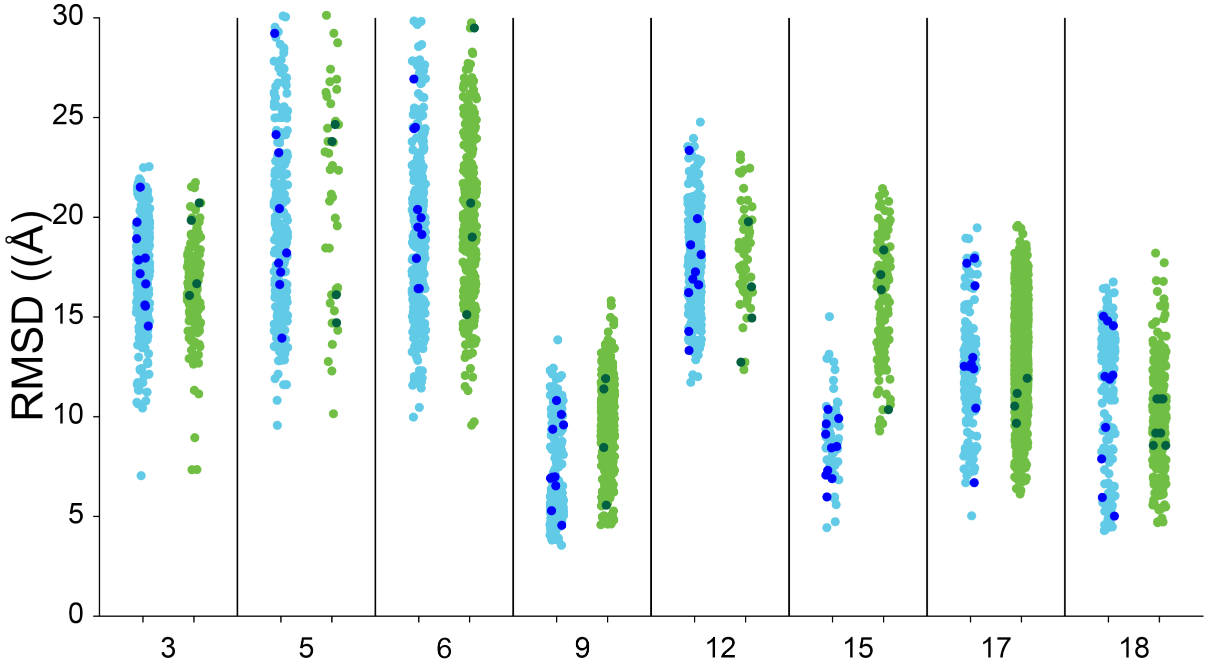

**Figure S5.** Related to Figure 3. A direct comparison of eight Puzzles run with base pair step sampling (i.e., standard FARFAR2; light blue points with dark blue cluster centers) with an identical protocol using fixed helical stems (preserving the scorefunction and fragment library; light green points with dark green cluster centers) confirms the utility of explicit sampling of helical flexibility.

**Table S1. Related to Figure 1.** Fragment assembly performance depends on multiple parameters optimized in this study. The performance of FARFAR2 (gauged by the lowest RMSD obtained from the 1% lowest energy models) was measured on the original FARNA benchmark to compare scoring functions, secondary structure input, fragment sets, and simulation guidance parameters. This study determined the ideal simulation parameters for use in the remaining larger-scale benchmarks.

| | | | | | | | | | | | | | | | | | | | | | | | | | no filters | -rounds 1 | -rounds 2 |
|---|---|---|---|---|---|---|---|---|---|---|---|---|---|---|---|---|---|---|---|---|---|---|---|---|---|---|---|
| Fragment library | Old | | | | | | | | | | | | New | | | | | | | | | | | | New | New | New |
| Mode of SS input | None | | | Fixed helix input | | | Secondary structure constraints | | | Base pair steps | | | None | | | Fixed helix input | | | Secondary structure constraints | | | Base pair steps | | | Base pair steps | Base pair steps | Base pair steps |
| Minimization scorefunction | n/a | hires | rna4 | n/a | hires | rna4 | n/a | hires | rna4 | n/a | hires | rna4 | n/a | hires | rna4 | n/a | hires | rna4 | n/a | hires | rna4 | n/a | hires | rna4 | rna4 | rna4 | rna4 |
| 157d | 1.385 | 0.797 | 0.513 | 1.363 | 0.763 | 0.556 | 1.288 | 1.288 | 1.288 | 2.003 | 1.258 | 0.941 | 0.839 | 0.653 | 0.457 | 1.470 | 0.786 | 0.476 | 1.065 | 1.065 | 1.065 | 1.644 | 0.947 | 0.618 | 1.018 | 1.651 | 0.944 |
| 1a4d | 5.266 | 6.185 | 5.352 | 3.370 | 2.884 | 2.556 | 5.581 | 7.052 | 3.786 | 3.381 | 2.823 | 3.101 | 4.785 | 4.412 | 3.782 | 3.863 | 3.035 | 3.240 | 6.818 | 4.651 | 3.732 | 3.751 | 2.737 | 3.036 | 3.341 | 3.503 | 3.495 |
| 1csl | 2.472 | 2.275 | 1.814 | 2.303 | 3.133 | 1.956 | 2.160 | 2.160 | 2.160 | 3.004 | 3.066 | 2.508 | 2.021 | 1.971 | 2.297 | 2.323 | 2.399 | 1.914 | 1.639 | 2.094 | 1.639 | 2.592 | 3.341 | 2.368 | 2.592 | 2.164 | 2.273 |
| 1dqf | 0.968 | 0.699 | 0.972 | 0.704 | 0.708 | 0.791 | 1.064 | 0.938 | 1.087 | 1.028 | 1.029 | 0.933 | 1.621 | 1.242 | 1.072 | 0.699 | 0.768 | 1.159 | 1.645 | 1.842 | 1.810 | 0.929 | 0.989 | 0.666 | 0.772 | 0.726 | 1.226 |
| 1esy | 3.141 | 2.334 | 2.217 | 1.866 | 1.743 | 1.543 | 2.788 | 3.170 | 3.134 | 2.269 | 2.087 | 1.995 | 2.459 | 2.459 | 2.459 | 1.965 | 1.925 | 1.637 | 2.770 | 2.834 | 2.770 | 2.048 | 2.130 | 1.842 | 2.059 | 1.989 | 1.871 |
| 1i9x | 1.866 | 1.801 | 1.819 | 2.504 | 3.264 | 2.937 | 1.887 | 1.947 | 1.887 | 2.425 | 2.753 | 2.462 | 1.675 | 1.715 | 1.677 | 2.643 | 3.382 | 2.978 | 2.853 | 2.857 | 2.293 | 2.285 | 2.320 | 1.558 | 2.315 | 2.361 | 3.396 |
| 1kd5 | 3.000 | 2.686 | 2.362 | 3.128 | 2.240 | 2.240 | 2.112 | 2.143 | 2.143 | 2.826 | 1.676 | 1.677 | 3.402 | 2.785 | 2.576 | 3.215 | 2.794 | 2.579 | 2.128 | 2.660 | 1.718 | 2.636 | 1.947 | 1.979 | 1.383 | 2.334 | 1.651 |
| 1kka | 3.785 | 3.546 | 3.546 | 5.233 | 5.263 | 5.210 | 3.603 | 3.688 | 3.482 | 3.262 | 3.111 | 3.377 | 3.888 | 3.739 | 3.486 | 5.155 | 5.090 | 5.223 | 3.616 | 3.531 | 3.413 | 3.215 | 3.331 | 3.252 | 3.693 | 3.579 | 3.493 |
| 1l2x | 2.415 | 2.935 | 9.981 | 2.255 | 2.302 | 2.410 | 2.715 | 2.885 | 2.715 | 2.270 | 2.996 | 2.664 | 2.261 | 2.360 | 8.813 | 2.293 | 2.196 | 2.276 | 2.449 | 2.922 | 2.922 | 2.434 | 2.483 | 2.763 | 3.414 | 2.727 | 2.499 |
| 1mhk | 7.539 | 6.591 | 6.639 | 5.770 | 4.458 | 5.079 | 6.451 | 4.264 | 4.310 | 3.668 | 3.848 | 4.305 | 5.114 | 4.921 | 5.204 | 5.473 | 5.195 | 4.835 | 6.988 | 6.409 | 6.505 | 5.809 | 4.556 | 4.735 | 4.780 | 6.025 | 4.716 |
| 1q9a | 4.156 | 3.645 | 4.082 | 4.013 | 4.111 | 4.173 | 3.954 | 3.869 | 3.954 | 4.085 | 2.178 | 3.248 | 4.184 | 4.144 | 4.184 | 3.989 | 2.298 | 4.024 | 4.295 | 4.448 | 4.448 | 4.008 | 3.234 | 3.217 | 1.581 | 4.067 | 4.091 |
| 1qwa | 3.378 | 3.293 | 3.733 | 2.747 | 2.928 | 3.136 | 3.291 | 3.456 | 3.344 | 2.869 | 3.480 | 4.005 | 3.613 | 3.345 | 2.911 | 2.928 | 2.652 | 2.863 | 3.328 | 3.397 | 3.328 | 2.358 | 2.729 | 3.375 | 2.967 | 2.793 | 3.840 |
| 1xjr | 12.32 | 9.913 | 9.919 | 7.659 | 6.623 | 6.694 | 8.115 | 7.520 | 6.531 | 7.127 | 6.352 | 6.270 | 11.00 | 9.254 | 8.636 | 5.848 | 4.950 | 7.057 | 8.375 | 7.141 | 7.130 | 7.178 | 7.241 | 7.012 | 7.197 | 7.815 | 5.855 |
| 255d | 1.354 | 1.282 | 1.296 | 1.556 | 1.146 | 1.078 | 1.221 | 1.234 | 1.234 | 1.488 | 1.517 | 0.982 | 1.339 | 1.173 | 1.249 | 1.237 | 1.070 | 1.160 | 1.208 | 1.208 | 1.208 | 1.162 | 1.155 | 1.201 | 1.135 | 1.016 | 1.290 |
| 283d | 1.651 | 1.780 | 1.702 | 4.086 | 1.163 | 1.305 | 1.566 | 1.469 | 1.714 | 1.746 | 1.865 | 1.717 | 1.723 | 1.443 | 2.357 | 4.993 | 1.167 | 1.226 | 1.619 | 1.516 | 1.879 | 2.548 | 1.572 | 1.337 | 1.623 | 1.684 | 1.361 |
| 28sp | 2.554 | 2.428 | 2.428 | 3.008 | 2.491 | 2.269 | 2.705 | 2.705 | 2.705 | 2.622 | 2.485 | 2.285 | 2.819 | 2.731 | 2.762 | 2.775 | 2.468 | 2.291 | 2.501 | 2.501 | 2.501 | 2.550 | 2.484 | 2.081 | 2.508 | 2.271 | 2.181 |
| 2a43 | 6.679 | 6.701 | 5.399 | 3.326 | 3.341 | 3.055 | 3.973 | 3.973 | 3.598 | 3.250 | 3.155 | 3.448 | 3.919 | 4.063 | 3.767 | 3.176 | 3.069 | 2.859 | 3.100 | 3.100 | 3.100 | 2.986 | 2.722 | 2.839 | 3.556 | 3.807 | 2.468 |
| 2f88 | 2.822 | 2.856 | 2.285 | 2.156 | 3.143 | 2.491 | 3.976 | 3.639 | 2.879 | 1.896 | 2.053 | 2.250 | 2.876 | 2.608 | 2.298 | 2.333 | 2.388 | 2.278 | 4.334 | 3.201 | 2.293 | 2.294 | 1.977 | 2.183 | 2.256 | 2.026 | 2.008 |

**Table S2. Related to Figure 1.** Comparison of fragment assembly performance using the modern scoring function on the 'motif-scale' benchmark set versus the performance of SWM on the same benchmark. More models were generated for the FARFAR version of the benchmark, but comparable computational time was required in each case. Unlike in the original SWM work, the relevant metric compared is the best RMSD sampled from among the 1% of models with the best energy.

| Benchmark Case | FARFAR 1% RMSD (A) | SWM 1% RMSD (A) | FARFAR Best of 5 Cluster RMSD | SWM Best of 5 Cluster RMSD |
|---|---|---|---|---|
| *Trans-Helix Loops* | | | | |
| 5P_j12_leadzyme | 2.56 | 0.82 | 2.56 | 1.22 |
| 5P_p1_m_box_riboswitch | 2.64 | 2.13 | 2.68 | 0.63 |
| 3P_j55a_group_I_intron | 0.27 | 1.72 | 0.42 | 0.40 |
| 5P_j55a_group_I_intron | 3.28 | 3.25 | 3.87 | 0.74 |
| hepatitis_C_virus_ires_IIa | 3.23 | 3.11 | 3.47 | 2.56 |
| j24_tpp_riboswitch | 1.38 | 1.22 | 3.51 | 0.72 |
| j23_group_II_intron | 1.77 | 1.11 | 2.57 | 0.55 |
| j31_glycine_riboswitch | 5.53 | 0.54 | 5.53 | 0.65 |
| l1_sam_II_riboswitch | 0.67 | 3.14 | 0.89 | 0.83 |
| l2_viral_rna_pseudoknot | 1.76 | 4.28 | 1.94 | 0.71 |
| 23s_rrna_44_49 | 1.20 | 0.71 | 1.23 | 1.25 |
| 23s_rrna_531_536 | 4.61 | 1.49 | 5.41 | 1.48 |
| 23s_rrna_2534_2540 | 1.97 | 7.28 | 2.90 | 6.94 |
| 23s_rrna_1976_1985 | 8.19 | 12.18 | 9.82 | 15.27 |
| 23s_rrna_2003_2012 | 7.54 | 9.85 | 9.85 | 9.02 |
| *Total cases: 15* | *6* | *9* | *3* | *12* |
| *Apical Loops* | | | | |
| gcaa_tetraloop | 1.36 | 1.18 | 1.55 | 1.14 |
| uucg_tetraloop | 2.57 | 2.29 | 2.03 | 1.14 |
| gagua_pentaloop | 0.75 | 2.94 | 0.76 | 1.10 |
| anticodon_phe | 1.30 | 2.82 | 2.21 | 2.24 |
| *Total cases: 4* | *2* | *2* | *2* | *2* |
| *Two-Way Junctions, Fixed* | | | | |
| puzzle1_alt_fixed | 1.98 | 0.409 | 2.29 | 1.72 |
| srp_domainIV_fixed | 0.50 | 1.034 | 0.69 | 0.90 |
| srl_fixed | 0.54 | 0.707 | 0.69 | 0.56 |
| kink_turn_fixed | 0.80 | 2.02 | 1.28 | 1.45 |
| j55a_P4P6_fixed | 1.78 | 3.20 | 1.86 | 0.55 |
| P5b_connect | 0.76 | 3.63 | 0.76 | 2.68 |

| | | | | |
|---|---|---|---|---|
| gg_mismatch_fixed | 1.87 | 0.32 | 2.03 | 0.32 |
| tandem_ga_imino_fixed | 1.07 | 0.86 | 1.21 | 0.87 |
| tandem_ga_sheared_fixed | 0.37 | 0.61 | 0.50 | 0.61 |
| hiv_rre_fixed | 0.36 | 0.87 | 0.45 | 0.35 |
| j44a_p4p6_fixed | 0.75 | 0.56 | 1.03 | 0.58 |
| just_tr_P4P6_fixed | 0.66 | 0.63 | 0.68 | 0.61 |
| r2_4x4_fixed | 0.72 | 1.44 | 1.20 | 1.49 |
| loopE_fixed | 0.51 | 1.74 | 0.72 | 1.74 |
| *Total cases: 14* | *9* | *5* | *6* | *8* |
| *Three-Way Junctions, Fixed* | | | | |
| hammerhead_3WJ_cat_fixed | 3.25 | 1.62 | 3.71 | 3.05 |
| hammerhead_3WJ_precat_fixed | 1.16 | 1.60 | 1.90 | 1.57 |
| VS_rbzm_P2P3P6_fixed | 0.76 | 0.74 | 1.09 | 0.57 |
| VS_rbzm_P3P4P5_fixed | 1.21 | 1.91 | 1.21 | 1.91 |
| hammerhead_3WJ_cat_OMC_fixed | 2.85 | 2.16 | 4.56 | 3.04 |
| *Total cases: 5* | *2* | *3* | *1* | *4* |
| *Tertiary Contacts, Fixed* | | | | |
| tl_tr_P4P6 | 0.55 | 1.07 | 0.55 | 0.64 |
| hammerhead_tert_fixed | 0.96 | 2.45 | 1.00 | 1.16 |
| kiss_add_fixed | 2.28 | 3.24 | 2.69 | 2.40 |
| kiss_add_L2_fixed | 0.52 | 1.50 | 0.69 | 0.71 |
| kiss_add_L3_fixed | 1.42 | 2.11 | 1.57 | 0.88 |
| puzzle18_zika_PK | 2.67 | 2.01 | 2.67 | 1.97 |
| gir1_p2.1p5_kiss_fixed | 1.19 | 1.94 | 1.62 | 1.99 |
| gir1_p2p9_gaaa_minor_fixed | 0.83 | 2.02 | 1.02 | 1.58 |
| t_loop_fixed | 0.78 | 2.27 | 0.78 | 0.95 |
| t_loop_modified_fixed | 2.37 | 1.24 | 0.98 | 1.33 |
| *Total cases: 10* | *8* | *2* | *7* | *3* |
| *Two-Way Junctions, Aligned* | | | | |
| gg_mismatch | 1.12 | 2.65 | 1.12 | 0.79 |
| tandem_ga_imino | 0.89 | 1.32 | 1.03 | 0.98 |
| tandem_ga_sheared | 0.49 | 1.06 | 0.61 | 0.75 |

| | | | | |
|---|---|---|---|---|
| hiv_rre | 1.78 | 3.59 | 1.94 | 2.12 |
| j44a_p4p6 | 1.01 | 4.75 | 1.56 | 1.59 |
| just_tr_P4P6 | 0.74 | 2.47 | 0.96 | 1.22 |
| cg_helix | 0.23 | 0.63 | 0.28 | 0.58 |
| puzzle1 | 0.91 | 3.26 | 0.84 | 0.96 |
| srp_domainIV | 0.94 | 2.88 | 1.01 | 1.26 |
| r2_4x4 | 1.84 | 3.34 | 1.84 | 1.74 |
| gagu_forcesyn_blockstackU | 4.65 | 5.64 | 4.87 | 4.49 |
| srl_free_bulgedG | 4.59 | 6.38 | 4.76 | 4.66 |
| j55a_P4P6_align | 0.85 | 2.81 | 1.04 | 2.04 |
| kink_turn_align | 0.79 | 3.01 | 0.97 | 2.07 |
| loopE | 0.91 | 5.28 | 1.75 | 2.00 |
| *Total cases: 15* | *15* | *0* | *10* | *5* |
| *Three-Way Junctions, Aligned* | | | | |
| hammerhead_3WJ_precat | 2.83 | 10.74 | 4.18 | 6.09 |
| VS_rbzm_P2P3P6_align | 0.73 | 1.29 | 0.84 | 1.13 |
| VS_rbzm_P3P4P5_align | 1.04 | 2.40 | 1.86 | 2.60 |
| hammerhead_3WJ_cat_OMC_align | 2.56 | 2.89 | 4.22 | 2.89 |
| puzzle18_zika_3WJ_extraminres | 2.52 | 4.36 | 2.99 | 2.42 |
| *Total cases: 5* | *5* | *0* | *3* | *2* |
| *Tertiary Contacts* | | | | |
| gaaa_minor_dock | 1.02 | 2.26 | 1.19 | 1.41 |
| gir1_p2.1p5_kiss | 1.48 | 3.25 | 2.01 | 2.75 |
| gir1_p2p9_gaaa_minor | 1.17 | 2.60 | 1.16 | 1.83 |
| tl_tr_P4P6_dock | 0.81 | 6.83 | 0.81 | 3.03 |
| kiss_add_PK_dock | 2.07 | 3.46 | 3.35 | 2.58 |
| t_loop_align | 2.02 | 4.16 | 2.02 | 3.20 |
| hammerhead_tert_align | 3.08 | 7.87 | 3.90 | 8.65 |
| t_loop_modified_align | 1.93 | 3.66 | 2.44 | 3.99 |
| *Total cases: 8* | *8* | *0* | *7* | *1* |
| *Non-Helix Embedded* | | | | |
| cg_helix_Zform | 5.00 | 10.76 | 5.00 | 1.75 |

| | | | | |
|---|---|---|---|---|
| g_quadruplex_fixed | 1.58 | 3.57 | 1.79 | 2.75 |
| g_quadruplex_inosine_fixed | 1.84 | 2.43 | 2.03 | 2.87 |
| bru_gag_tetraplex | 3.30 | 2.78 | 2.70 | 3.42 |
| parallel_AA | 0.97 | 1.22 | 1.16 | 1.41 |
| bulged_tetraplex | 4.95 | 7.67 | 5.04 | 7.37 |
| *Total cases: 6* | *5* | *1* | *5* | *1* |
| *Overall: 82* | *60* | *22* | *44* | *38* |

**Table S3. Related to Table 1.** Exact inputs (sequence and RNA templates) used in modeling for each RNA-puzzle problem.

| Puzzle | Secondary Structure | Inputs |
|---|---|---|
| 1 | ccgccgcgccaugccuguggcgg,ccgccgcgccaugccuguggcgg<br>(((((((..((((...(((((((,)))))))..))))...))))))) | Just secondary structure |
| 2 | ccggaggaacuacugccggcagccuccggaggaacuacugccggcagccuccggaggaac<br>uacugccggcagccuccggaggaacuacugccggcagccu<br>{{{{(((.....(((aaaa))))))<<<<(((.....(((}}}}))))))[[[[(((...<br>..(((>>>>))))))aaaa(((.....(((]]]])))))) | Four of eight chains were provided with puzzle specification. Three 'corners' modeled as HCV IRES loops. |
| 3 | cucuggagagaaccguuuaaucggucgccgaaggagcaagcucugcgcauaugcagagug<br>aaacucucaggcaaaaggacagag<br>(((((......((((......)))))(((...((((...((((((((....)))))))).<br>...)))...))).......))))) | Just secondary structure |
| 4 | ggcuuaucaagagagguggagggacuggcccgaugaaacccggcaaccacuagucuagcg<br>ucagcuucggcugacgcuaggcuaguggugccaauuccugcagcggaaacguugaaagau<br>gagcca<br>................................................(((((.((((((<br>(((((....))))))))))).)))))..................................<br>...... | 3IQP, the template at the time, provides residues A:1-47 A:88-126 |
| 5 | gguuggguugggaaguaucauggcuaaucaccaugaugcaaucggguugaacacuuaauu<br>ggguuaaaacggugggggacgaucccguaacauccguccuaacggcgacagacugcacgg<br>cccugccucuuagguguguccaaugaacagucguuccgaaaggaagcauccgguauccca<br>agacaauc<br>(((((..(((((..(((((((((...[[[.)))))))))..((((.(((.((((......<br>(.........((..(((...]]])))).......))........................<br>..)............)))).)))..........((((....))))....))))...))))<br>)..))))) | Template structure 3BO3 (group I intron) used in original Das lab modeling for A:62-70 A:96-122 A:148-153; A:55-60 A:124-129 A:132-135; they are thus omitted from the secondary structure. |
| 6 | cggcaggugcucccgac,gucgggaguuaaaagggaagccggugcaaguccggcacgguc<br>ccgccacugugacggggagucgccccucgggaugugccacuggcc,ggccgggaaggcgg<br>aggggcggcgaggauccggagucaggaaaccugccugccg<br>((((((((.(((.(((.,.)).)))).....((....(((((.......)))))[[((..<br>...)).(((...(((...(((((((.((.......(((.((((((,))))))...))).)<br>).)))))))......)))..]])))).....))))))))) | Template structures 2YIE (FMN aptamer) and 2GIS (SAM riboswitch) used in original Das lab modeling provided A:5-8 A:32-35 A:37-39 A:158-164; A:81-83 A:138-143; A:41-61 A:67-75 A:147-149. |
| 7 | gcgcugugucgcaaucugcgaagggcgucgucggcccaagcgguaguaagcagggaacuc<br>accuccaaugaaacacauugucguagcaguugacuacuguuaugugauugguagaggcua<br>agugacgguauuggcguaagccaauaccgcagcacagcacaagcccgcuugcgagauuac<br>agcgc<br>((((((((((.....((((...((((.......))))..))))..((((((.(((..(((<br>(((..(((((.....)))))..(((((((.......)))))))......))).)))(((.<br>.(((.((((((((((....))))))))))...)))))).....)))))))))..))).))<br>))))) | Just secondary structure. |
| 8 | ggaucacgagggggagaccccggcaaccugggacggacacccaaggugcucacaccggag<br>acgguggauccggcccgagagggcaacgaaguccgu<br>(((((....((((....))))(((..((((((.[[[[..))).))).)))..(((((...<br>.)))))))))).((((....)))).......]]]]. | Template structures from original Das lab modeling for SAM binding site using 2YGH, as described previously. |

| | | |
|---|---|---|
| 9 | ggacacugaugaucgcguggauauggcacgcauugaauuguuggacaccguaaauguccu aacacguguccc<br>((((((......(.(((((.....[[))))))..........(((((]].....))))).<br>.....)))))) | Template structures from original Das lab modeling: kissing loop from 2XNW; T-loop from the tRNA Phe structure 3L0U. |
| 10 | ugcgaugagaagaagaguauuaaggauuuacuaugauuagcgacucuaggauagugaaag cuagaggauaguaaccuuaagaaggcacuucgagca,aguaguucagugguagaacacca ccuugccaagguggggucgcggguucgaaucccgucu<br>(((.....(((.......((((((..(((((((.........((((((...........<br>))))))).)))))))))))))))...[[[.)))))..))),<br>((..((((....{..)))).(((((..]]]..))))).....(((((..}....))))))<br>) | Template structures from original Das lab modeling: a kink-turn from 2GIS provided A:1-10, A:90-96; a loopE motif from 5S rRNA 354D furnished A:16-20, A:80-83; a double T-loop motif from 4JRC supplied A:37-43, A:53-60; and part of a tRNA from 2K4C delivered B:6-18, B:20-3,0 B:38-66. |
| 11 | gggaucugucaccccauugaucgccuucgggcugaucuggcuggcuaggcggucccc<br>((((((((((.(((((..((((((((....))).)))))))..))...)))))))))) | Just secondary structure |
| 12 | gaucgcugaacccga,aggggcgggggacccag,ggggcgaaucucuuccgaaaggaaga guaggguuacuccuucgacccgagcccgucagcuaaccucgcaagcguccgaaggagaa<br>...((((...(((..,.)))((((((....(..,(((((....(((((((....))))))<br>)..(((((.[[[[[[[))))))..))))..).)....)))))).))))...]]]]]]].. | Just secondary structure |
| 13 | gggucgugacuggcgaacaggugggaaaccaccggggagcgacccc,gccgcccgccuggg c<br>(((((((..(([[[[....(((((....))))).))..))))))),(((...]]]]..))<br>) | Just secondary structure |
| 14b | cguugacccaggaaacugggcggaaguaaggcccauugcacuccgggccugaagcaacgc g<br>(((((.(((((....)))))........((((((..........))))))....))))).<br>. | Just secondary structure (original puzzle omits U1A loop in RMSD calculations) |
| 14f | cguuggcccaggaaacugggu,aguaaggcccauugcacuccgggccugaagcaacgcu<br>(((((((((((....)))))),....((((((..........))))))....))))).. | Just secondary structure (original puzzle omits U1A loop in RMSD calculations) |
| 15 | ggguacuuaagcccacugaugagucgcugggaugcgacgaaacgccca,gggcgucuggg caguaccca<br>..........((((((....).(((((......)))))...((((((.,)))))).))))<br>)........ | Just secondary structure |
| 17 | cgugguuagggccacguuaaauaguugcuuaagcccuaagcguugau,aucaggugcaa<br>((((([[[[[[))))))........((((.....]]]]]]....(((.,.)))...)))) | Just secondary structure |
| 18 | gggucaggccggcgaaagucgccacaguuuggggaaagcugugcagccuguaacccccccc acgaaagugggg<br>....(((((((((....)))).((((((..[[[[...))))))..)))))...]]]](((<br>((....))))) | Template structures from original Das lab modeling from 4PQV. |
| 19 | gcagggcaaggcccagucccgugcaagccgggaccgcccc,ggggcgcggcgcucauucc ugc<br>(((((...(((((..((((((.......))))))((((((,)))))).))).))....))<br>))) | Template structures from original Das lab modeling: threaded T-loop from 1B23, with intercalating A:8 modeled in. (Other suspected intercalators like A:7 and B:15 score strictly worse.) |
| 20 | acccgcaaggccgacggc,gccgccgcuggugcaaguccagccacgcuucggcgugggcg cucaugggu | Template structures from original Das lab modeling: threaded T-loop from puzzle 19 (5T5A), renumbered. |

| | | |
|---|---|---|
| | ((((...((((((.((((,)))).)(((((.......)))))(((((....))))))))).<br>))...)))) | |
| 21 | ccggacgaggugcgccguacccggucaggacaagacggcgc<br>[[[[.......(((((((..]]]]...........))))))) | Just secondary structure, no guanidinium. |

**Table S4. Related to Table 1.** Exact inputs (sequence and RNA templates) used in modeling for each RNA-puzzle problem.

| Blind challnege | Secondary Structure | Inputs |
|---|---|---|
| *F. nucleatum* glycine riboswitch | ucggaugaagauaugaggagagauuucauuuuaaugaaacaccgaagaag<br>uaaaucuuucagguaaaaaggacucauauuggacgaaccucuggagagcu<br>uaucuaagagauaacaccgaaggagcaaagcuaauuuuagccuaaacucu<br>cagguaaaaggacggag<br>(((......((((((((......((((((....)))))).(((...((((<br>.....))))..)))........))))))))...)))..(((((......(<br>(((((...)))))).(((....(((....((((....)))).....))).<br>..))).......))))) | 3P49 furnished residues A:19-29 A:34-71 A:92-164. |
| *V. cholerae* glycine riboswitch | uccguugaagacugcaggagagugguuguuaaccagauuuuaacaucuga<br>gccaaauaacccgccgaagaaguaaaucuuucaggugcauuauucuuagc<br>cauauauuggcaacgaauaagcgaggacuguaguuggaggaaccucugga<br>gagaaccguuuaaucggucgccgaaggagcaagcucugcgcauaugcaga<br>gugaaacucucaggcaaaaggacagagga<br>[[[......((((((((......(((((((...(((((......))))).<br>....))))))).(((....(((.....)))...)))((.((((((...((<br>((.....))))...))))))))).....))))))))...]]]..(((((..<br>..(.((((......)))))(((...((((...(((((((......)))))<br>))....))).).)))......))))).. | Threaded template from lowest energy model of *F. nucleatum* glycine riboswitch modeling above: residues A:1-29 A:34-64 A:68-91 become A:1-29 A:56-86 A:125-148. |
| SAM-IV riboswitch | ggucaugagugccagcgucaagccccggcuugcuggccggcaacccucca<br>accgcggugggggugccccgggugaugaccagguugaguagccgugacggc<br>uacgcggcaagcgcgguc<br>((((....(.((((((....((.[[[[[)).)))))))(((..(((.((.<br>.{{{.))..))).)))]]]]]....))))..((((.((((((.....)))<br>))).)))).....}}}... | SAM binding site from 2YGH provided A:1-9 A:38-43 A:63-66 A:76-79, with SAM as B:120 |
| Spinach-TTR-3 | ggcgauauggauagaaggacggguccccauauugcgagaaacguuauacgg<br>aaccguacgaaggaaacuucguacucaguacguauaacggagcaauaugg<br>uugaguagagugugagcuauccuaagucgc<br>.(((((.(((((((.....(...(((((((((((......(((((((((.<br>....((((((((....))))))))......)))))))))..)))))))))<br>..))..........)..))))))..)))))) | Tetraloop/receptor from 1GID for residues A:62-67, A:120-127, A:5-11; DFHBI-binding G-quadruplex from 4TS2 for residues A:17-18 A:21-24 A:101 A:104-105 A:108 A:110 A:112 |
| VA RNA I | ggaccucgcaaggguaucauggcggacgaccgggguucgaaccccggauc<br>cggccguccgccgugauccaugcgguuaccgccccgcgugucgaacccagg<br>ugugcgaggucc<br>((((((((((.((((.((((((((((((.(((((.((........)).))<br>))).))))))))))))..(((((((..[[[..)))))))....)))).]]<br>].)))))))))) | Just secondary structure |
